# Building programmable multicompartment artificial cells incorporating remotely activated protein channels using microfluidics and acoustic levitation

**DOI:** 10.1101/2022.01.13.476178

**Authors:** Jin Li, William D. Jamieson, Pantelitsa Dimitriou, Wen Xu, Paul Rohde, Boris Martinac, Matthew Baker, Bruce W. Drinkwater, Oliver K. Castell, David A. Barrow

## Abstract

Intracellular compartments are functional units that support the metabolic processes within living cells, through spatiotemporal regulation of chemical reactions and biological processes. Consequently, as a step forward in the bottom-up creation of artificial cells, building analogous intracellular architectures is essential for the expansion of cell-mimicking functionality. Herein, we report the development of a droplet laboratory platform to engineer customised complex emulsion droplets as a multicompartment artificial cell chassis, using multiphase microfluidics and acoustic levitation. Such levitated constructs provide free-standing, dynamic, definable droplet networks for the encapsulation and organisation of chemical species. Equally, they can be remotely operated with pneumatic, heating, and magnetic elements for post-processing, including the incorporation of membrane proteins; alpha-hemolysin; and large-conductance mechanosensitive channel (MscL) and their activation. The assembly of droplet networks is three-dimensionally patterned with fluidic inputs configurations determining droplet contents and connectivity. Whilst acoustic manipulation can be harnessed to reconfigure the droplet network *in situ*. In addition, a mechanosensitive channel, MscL, can be repeatedly activated and deactivated in the levitated artificial cell by the application of acoustic and magnetic fields to modulate membrane tension on demand. This offers possibilities beyond one-time chemically mediated activation to provide repeated, non-contact control of membrane protein function. Collectively, this will expand our capability to program and operate increasingly sophisticated artificial cells as life-like materials.

## Main text

Through evolution, eukaryotic cells have exploited intracellular compartmentalisation to increase the efficiency and range of functional complexity in cellular metabolism [1]. There is wide interest in harnessing such principles in the creation of life-like synthetic materials. The bottom-up construction of artificial cells aims to harness and imitate some of these key cellular functions, with *de novo* structures [2,3]. Protocells mimic possible primitive cells, and have been routinely fabricated by forming small, singularly compartmentalised droplets with relatively simple infrastructure [4, 5], usually built through the macromolecular assembly of lipids or other amphiphiles [6, 7]. Such basic models can encapsulate different reagents, representing minimal ‘cellular’ systems for cell-free studies, including those of membrane properties [8], chemically mediated communication [9, 10, 11], and the manipulation of genetic information [12]. State-of-art protocell materials can interact with living cells [13] for potential biotechnological applications, such as immunogenicity enhancement [14], organoid formation [15] or blood vessel vasodilation [16].

To fabricate more complex artificial cells, a key objective is to develop new functionalities that are underpinned by intracellular compartmentalised architectures. Such structures can be formed by the encapsulation of lipid-bounded aqueous droplets and biomolecule complexes, within a host (envelope) droplet [17, 18], where each droplet may contain different biochemical species. Recent work has demonstrated that organelle-like components can work as functional units to process molecular signals [19, 20], regulate sequential reactions [21, 22, 23], and may be used for energy harvesting [24] within artificial cells. Furthermore, compartmentalisation has been harnessed in similar ways in the packing and patterning of individual protocells to construct more complex, functional, tissue-like materials, incorporating protein channels [25], DNA sequences [26], and functional hydrogels [27]. These works indicate that structural complexity can display a range of emergent properties defined by the contents and connectivity of constituent components in biomimetic materials. However, to date, few efforts have focused on the permutations of compartments within artificial cells. This is, in part, due to the fact that higher-order emulsification processes for the creation of multi-compartmental structures, with the controlled distribution of (bio)chemical reagents within such compartments, remains a very significant challenge. Most work to date has relied largely on the manual juxtaposition, or 3D-printing, of multiple droplets to create tissue-like materials [28, 29, 30]. Meanwhile, as the strategy of synthetic biology shifts from exploratory research to systematic quantified study, precision engineering will have a key role to play for the definable, and repeatable construction of artificial cells [31, 32]. Droplet microfluidics provides the ability to control droplet and vesicle formation and has become an invaluable tool in the fabrication of emulsion-based protocells [33,34,35]. Acoustic manipulation is a contactless, and non-invasive tool [36] that has been applied to protocell formation in microfluidic environments [37]. However, to date the full range of new opportunities for artificial cell construction, manipulation and operation provided by the combination of these techniques has yet to be fully realised. Current technological bottlenecks have limited the architectural complexity of precisely formed multicompartment artificial cells more generally. However, the application of programmable droplet microfluidics alongside acoustic manipulation and processing represents a novel route to affording increased control over compartmental organisation, connectivity, and communication, alongside opportunities for spatial and temporal control of protein function. These facets will pave the way for more routine production of artificial cell materials with the ability to engineer more complex and life-like characteristics, more akin to their biological counterparts.

Here, we report the development of a droplet laboratory platform for the creation, manipulation, control, and measurement of artificial cells with distributed cores (ACDC), using controllable droplet microfluidics and acoustic levitation (Fig.1 A-1 & A-2, Fig. S1). Membrane proteins can be incorporated in the levitated ACDC droplets and their functions remotely controlled *in situ* by the acoustic levitator. The ACDC droplets are defined as complex emulsion droplet materials with organised chemical encapsulations in hierarchical structures. These structures aim to mimic eukaryotic cell compartmentalisation and their endowed functionalities. In this work, ACDC droplets takes the form of a multisome model [30] suspended in air. Complex emulsions are constructed from microfluidically-formed, lipid-coated water droplets, here termed “cores”, where protein mediated communication can take place between bilayer segregated cores of the network. Once an ACDC droplet is dispensed from the microfluidic manifold, it can be trapped within the nodes (zones) of a vertically orientated acoustic standing wave, generated by a multi-emitter, single-axis acoustic levitator operating at 40KHz (Fig.1B). The droplet trapping trajectories depend on the droplet release positions (Fig.1C & Fig.S2), and the levitated droplets typically adopt an ellipsoid shape that can be controlled by the power of the acoustic transducer array (Fig.S3). The acoustic streaming effect [38] generates convective flows within the levitated droplet, shown by numerical simulations (Fig.1D), and these fluid dynamics may facilitate enhancement of mass transport in each phase of the emulsion droplet along the streamlines. We observed lipid bilayer formation within the levitated ACDC droplet, where lipid monolayer-coated water cores contact each other (Fig.1E). As high acoustic impedance exists at the air-liquid interface, the amount of acoustic radiation energy penetrating into the droplet is limited, and lipid bilayers remain robust against and do not fail or become porous. As the total volume of the internal core network increases, the assembled droplet interface bilayer networks rotate around the vertical axis (SI video 1) due to the induced acoustic streaming. This induced motion can be affected by external perturbation; for example, by the application of external air currents, or other asymmetric forces exerted on the ACDC droplet, such as applied magnetic fields. Without disruption of the acoustic standing wave field, post processing of levitated ACDC droplets *in situ* is possible, such as by the addition or removal of cores whilst the ACDC remains levitated in the trapping zone. Multiple ACDC droplets can be levitated and aligned at different nodes of a single standing wave field for parallel operations (Fig.1F). By using acoustic levitation for the control of artificial cells we demonstrate in this study that this system provides the ability to reproducibly process, activate, manipulate, and observe multicompartment artificial cells by contactless methods, immediately following their microfluidic production. We demonstrate control of droplet patterning within the ACDC capsule, that lipid bilayers can be broken and reformed on demand, and that internal core networks can be dynamically reorganised in situ. We demonstrate the ability to remotely activate the mechanosensitive channel of large conductance (MscL) on demand, enabling the switching on, and off, of artificial cell activity. Collectively this will enable the harnessing of more complex functionality in artificial cells, providing additional control of communication networks in space and time and moving away from single time activation.

**Figure 1.**
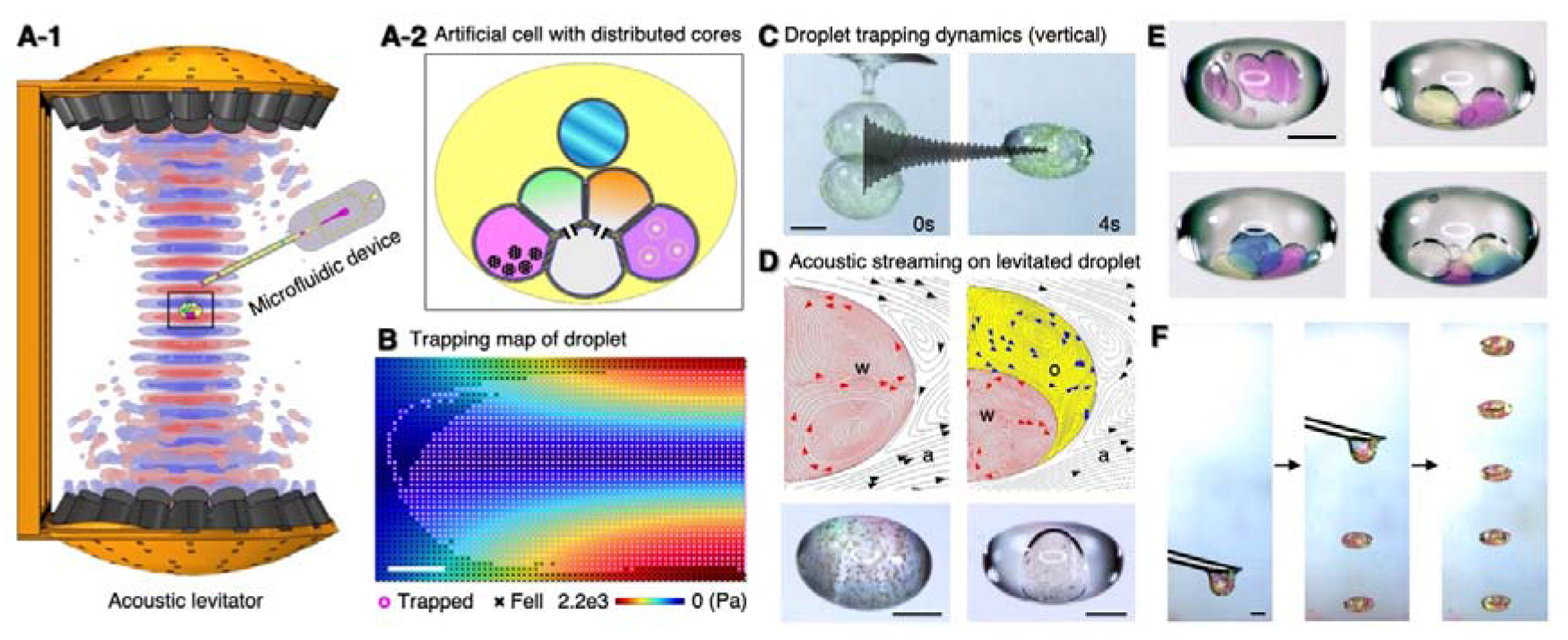
Droplet laboratory platform. **A-1**, schematic of the acoustic levitator platform setup. Multiple acoustic standing waves (red and blue), generated by the acoustic levitator, can trap microfluidically formed artificial cells with distributed cores (ACDC), at nodal positions. Photographs of full experimental apparatus are available in Fig. S1. **A-2**, schematic of patterned ACDC droplets encapsulating multiple reagents within the core networks. **B**. Simulation results of 2 mm diameter droplet trapping in air. The pink circle (droplet levitated) and the black cross (droplet fell) symbols represent the outcome of simulated droplet release positions in the acoustic field (rainbow colour map indicates standing wave pressure profile (symmetric half)). **C**. Droplet trapping dynamics: An ACDC droplet dispensed from the microfluidic device into a standing wave field oscillates vertically in the standing wave node before dampening and assuming a stable position over a period of 4 seconds. Additional data on droplet release positions are available in Fig. S2. **D**. Top - Simulation and experimental results of the convective flows within levitated water (w) droplet in air (a) (left) and water in oil (o) double emulsion droplets in air (a) (right). Bottom - Water phases containing hydrophilic magnetic microparticles (dia. 1 μm) and minor aggregates (brown), remain suspended and circulate with the fluidic convection, without significant clustering. **E**. Manually adding multiple water droplets to a levitated oil droplet with a micropipette. From top left to bottom right, pink, green, blue and clear droplets. **F**. Deposition of multiple microfluidically formed ACDC droplets at sequential nodes within the acoustic standing wave field. Scale bars = 1 mm.

### Control of ACDC droplet patterning

In our previous work, precursor ACDC droplets were developed as a free-standing artificial cell model that can harness membrane mimetic chemistry via encapsulated droplet interface bilayers [39]. 3D microfluidics enables the construction of polarised and directional ACDC droplets with patterned, multi-layered shells encapsulating functional nanomaterials [40]. Herein, we present a microfluidic approach to fabricate ACDC droplets with three-dimensionally organised interior core networks. A multi-layered, 3D-printed droplet forming junction was devised (Fig. S4), the design of which compensates for the resolution limitations of fused filament 3D printing, to form uniform, size-controlled (minimum 20 μm diameter - Fig. S4), lipid-coated aqueous droplets (cores) in a continuous oil phase flow. An array of these junctions is configured within a fluidic circuit to generate multiple types of cores in parallel (Fig.2A). Meanwhile, programmed inflow pressure profiles are applied to regulate the droplet formation on-demand, and produce droplet core sequences which template the order of reagent encapsulation (Fig.2B). Such 1D linear core sequences lead to the packing and self-organisation of patterned 3D core networks during the ACDC droplet formation and levitation (Fig 1C & SI video 1). The final configuration and connectivity of core networks is collectively influenced by the droplet order, microfluidic inlet flow profiles, microfluidic circuit configuration, and droplet trapping in the acoustic field. In this way, a target core containing specified reagents can be allocated to different regions of the core network for definable compartmentalisation within the ACDC droplet (Fig. 2C). This can be used for structuring the intracellular chemical organisation that is segregated by lipid bilayers with approximate control of the core’s coordinates in three dimensions. Such effort may be used to control compartment connectivity and separation and thus program sequential reactions within multicompartment artificial cells.

**Fig 2.**
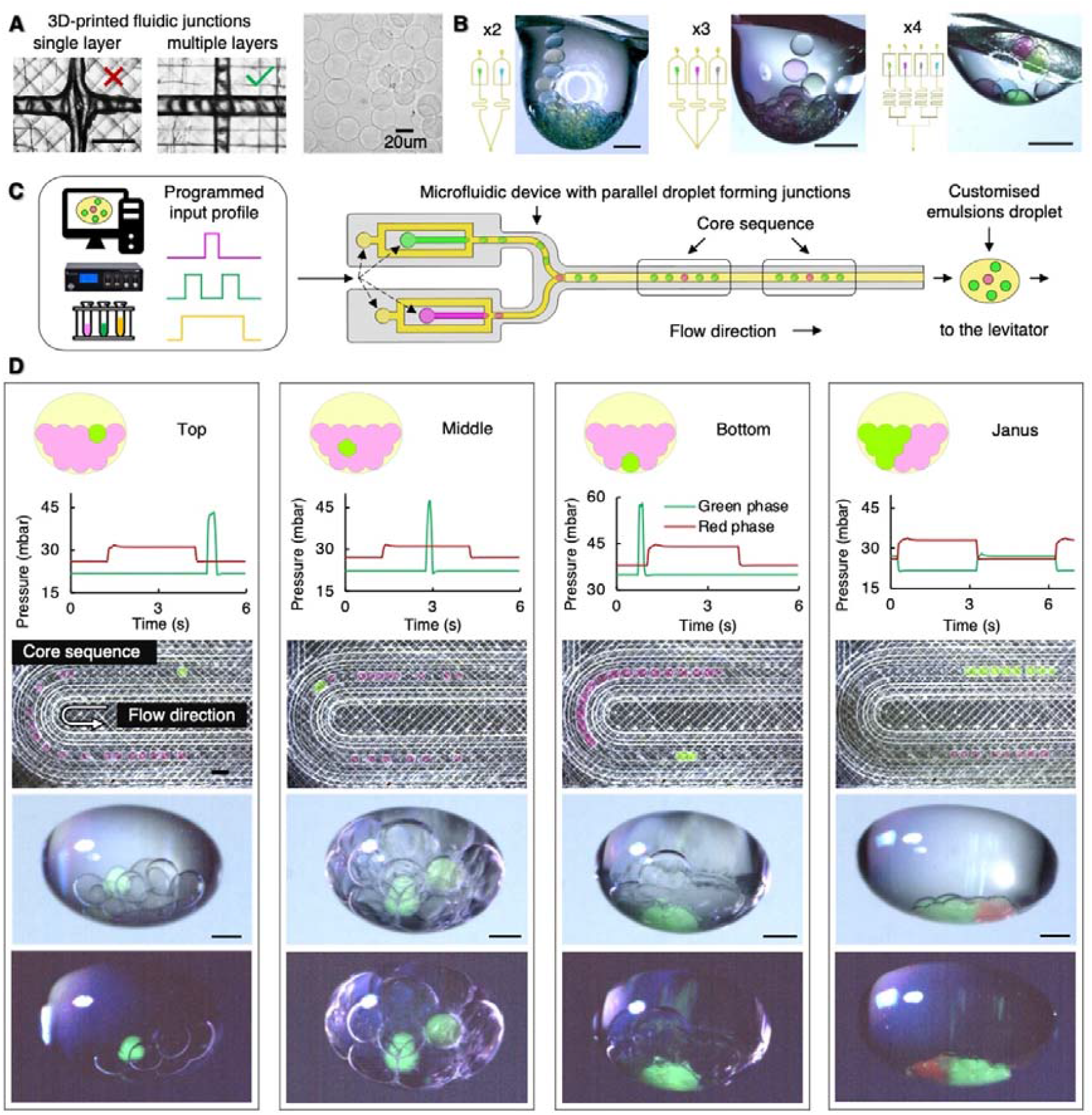
Microfluidic control of ACDC structural organisation. **A**. Multi-layered 3D printing minimises capillary junction size, improving droplet generation fidelity, in comparison to single layered devices (left), see Fig S4 for further details. Uniform, cell-sized, alginate microgels can be formed using 120 μm wide, 3D-printed multi-layered junctions (right) (flow rates: dispersed phase: 1 ml hr^-1^ and continuous phase: 20 ml hr^-1.^ Additional fluidic designs and droplet size data available in Fig. S4). **B**. The addition and combination of parallel fluidic junctions enables formation of multiple different types of water droplets within an oil compartment. Increasing complexity is illustrated left to right with cores of two, three and four different chemical identities. **C**. Programming of fluid inlet flows can be used to generate droplet sequences of defined order and spacing for the creation of customised emulsion droplets. Programmed input profiles regulate droplet formation at each droplet forming junction, on-demand, determining core sequences and therefore providing the chemical encapsulation template for the ACDC construct. Droplet order determines final 3D spatial arrangement of the encapsulated cores. **D**. Examples of patterned, levitated ACDC constructs. The fluidic input flow profile determines the core sequence and packing order in the ACDC construct. From left to right, a specific core (green) is directed to the top, middle and bottom level of ACDC constructs, and the formation of Janus networks comprised of red and green cores. Lipid bilayers segregate cores of the internal droplet network. Scale bars = 500 μm.

### Contactless operation of levitated ACDC droplet

Microfluidically formed, lipid membrane segregated, complex emulsion droplets provide metastable configurations for chemical organisation and connectivity. Meanwhile, in comparison to the batch processing methods used for single compartment protocells in aqueous environments, we present a comprehensive approach to operate, process and activate individual levitated ACDC droplets and their sub-compartments using a range of contactless methods for physically chemical processing (Fig. 3A, SI Video 2). This may enhance the breadth of multicompartment artificial cell functionalities, enabling chemical modulations of artificial cells [41].

**Fig 3.**
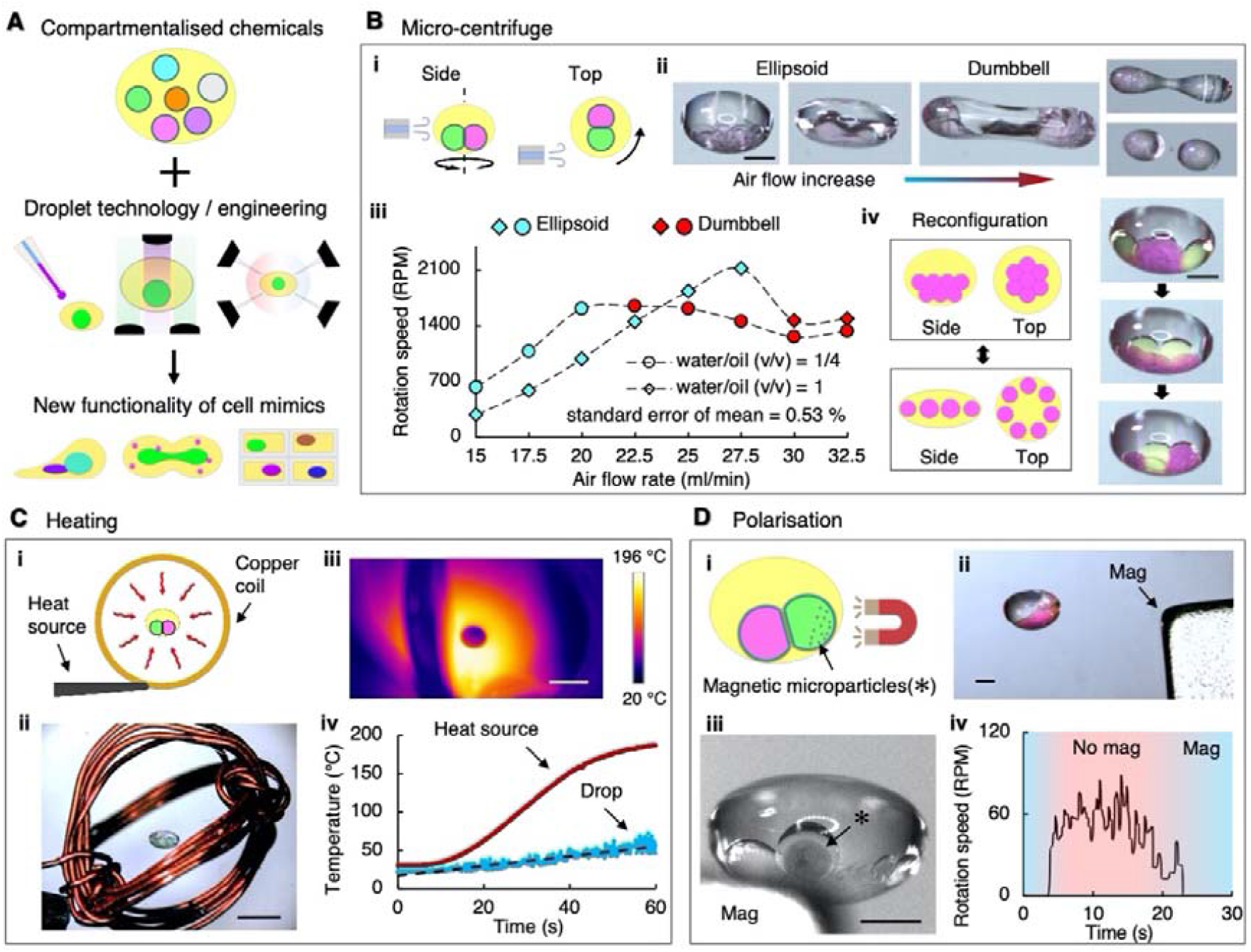
Multiple operations on levitated ACDC droplets. **A**. The combination of controlled chemical compartmentalisation, with subsequent droplet processing and manipulation, as methods to energise the droplet structures, can be used to impart new function on the resulting materials. **B**. Pneumatic operations for the spinning of levitated ACDC droplets. (i) A radially tangential air flow to the ACDC droplet rotates the levitated construct. (ii) As the air flow is increased, the droplet rotational speed increases, and the droplet becomes elongated before splitting into two daughter droplets, each with interior core network. (iii)The spinning rate can be precisely controlled by the air flow rate (inset graph: n=30 for each data point, measured from a 20s spinning interval). (iv-left column) At high rotational speeds the droplet network disassembles and lipid bilayers are separated between adjoining cores. Spin-stop cycles can be used to reconfigure the internal core network, detaching and re-assembled bilayer connections. (iv-right column) This is depicted for disassembly and reassembly of a core network (green and pink cores) being reconfigured to separate and connect different cores. Scale bars = 1 mm. **C**. Thermal operations for the heating of levitated ACDC droplets. A copper wire coil was used to induce thermal energy within the levitated droplet through thermal irradiation (i & ii). The droplet can be heated up from room temperature to 50°C in 1 minute with gradual heating (iii & iv). Scale bars = 5 mm. **D**. Magnetic operations for the manipulation of levitated ACDC droplets (i&ii). A levitated ACDC droplet containing a core with encapsulated magnetic particles (iii), can be manipulated by external magnets, providing droplet positional and orientational control (iv). The star symbol (*) indicates the position of the magnetic particles, and their gathering at the core perimieter on application of an external magnetic field. Scale bars = 1 mm.

The first demonstration utilises a pneumatic device such that the local air flow controls the rotation of ACDC droplets (Fig. 3B). As shown in the figure, an air nozzle is carefully controlled to tangentially apply a thin air stream to the equator of the levitated droplet (Fig 3B-i). This causes the droplet to spin about its central vertical axis whilst remaining trapped in a pressure node of the levitator. With increasing airflow, the morphology of the ACDC droplet can be changed from ellipsoid to a dumbbell shape, and eventually breaking it up into two daughter droplets, splitting as a result of droplet thinning at the central waist (Fig 3B-ii). The shape and angular velocity of spinning ACDC droplets can be precisely controlled by the air inflow rate and stably maintained up to 35 revolutions per second as shown in Fig. 3B-iii (n=30 for each data point, average standard error of mean = 0.53%). Whilst spinning, the surface area of interior droplet interface bilayers is minimised, and the droplet contact angle is increased until the membrane leaflets separate and the droplets fully detach. This is a consequence of the centrifugal force resulting in the dense water core moving radially outwards in the less dense oil phase (Fig S5). For a given core network, this detachment depends on the spinning angular velocity and droplet size, as well as the oil phase composition and the acoustic transducers power (Fig S5). On cessation of air flow, the spinning of the ACDC droplet slows down, and the separated cores reassemble, again forming lipid bilayers between contacting cores. New core networks may be assembled in this way, with the volumes and densities of the cores determining their release order and reassembly. This bilayer disassembly-assembly process can be used to reconfigure the core network patterns within the ACDC droplet, changing the packing order. For instance, Fig 3B-iv shows reconfiguration by this method of an internal core network comprised of green and red droplets, creating different patterns of core-core connectivity, following spin-based separation and reassembly. No core coalescence or dye mixing was observed indicating that the bilayer membranes were intact during the spinning, separation and reformation processes. Since membrane protein channels can be inserted into the bilayers of core networks, this rearrangement of connectivity can determine chemical communication and consequently function. This represents a method to reconfigure an artificial cell on demand, for the activation, or selection, of different functionalities dependent upon the organisation of the cores. Despite the complexity of this structuring mechanism, more robust protocols of core reconfiguration can be established from the mathematical modelling of sphere packing and acoustic assembly [42, 43].

A levitated ACDC droplet can also be energised and operated remotely utilising convective heating, photo-irradiation, or magnetic manipulation. Fig. 3C-i shows the heating of droplets via thermal irradiation. Copper coils (Fig. 3C-ii) were crafted to transfer thermal energy to the levitated droplet by conduction without disruption of the acoustic standing wave. In this way the droplet can be efficiently heated up from ambient temperature to ~50 °C in one minute (Fig. 3C-iii&iv). While the temperature gradually increased, the levitated droplet started to agitate and adopt a discoid architecture, possibly due to changes in the air, oil, and water properties (e.g. density, viscosity, surface tension) in response to the heating. The disk shape of the heated droplets can be tuned back to ellipsoid by gradually reducing the power input to the ultrasonic transducers, and vice versa. This heating operation can be applied to initiate thermo-responsive reactions within the ACDC droplet. Fig. 3D-i shows the employment of magnetic manipulation and ability to control the orientation of a levitated ACDC droplet. One specific compartment of the droplet network containing magnetic microparticles is attracted to an externally applied magnetic field, orientating the ACDC droplet in the levitator trap (Fig. 3D-ii&iii). Hence, the whole core network is in turn re-orientated via the membrane connections. Under the applied magnetic field, the ACDC droplet becomes slightly offset from the original equilibrium position, given the new balance of both magnetic field and acoustic field forces. In this way, the levitated ACDC droplet can be prevented from passive rotation or spinning induced by other operations or disturbances (Fig. 3D-iv). The orientation of the core networks can be precisely controlled (e.g. for observation) by the positioning of the external magnet in the system. In a further example of contactless processing, light-initiated reactions, such as the free-radical polymerisation of microfluidically formed poly(ethylene glycol) diacrylate (PEGDA) droplet containing an internal oil core, may be implemented in a levitated droplet (Figure S6). The heat generated by the exothermic reaction and its subsequent dissipation can be measured *in situ*. This indicates the possibility to fabricate free-standing, hierarchical materials with soft interiors and rigid shells, such that the processed ACDC droplets can be stored or transported to other environments.

### Reconstitution of protein channels in levitated ACDC droplets

Establishing controllable chemo-mediated communication between compartments, provides a means to govern biochemical processes within artificial cells. In a lipid-based model, intracellular communications can be controlled by the incorporation of membrane protein channels and pores to selectively permeabilise segregating membranes. Herein, first we demonstrate that the lipid bilayers of the ACDC droplet remain stable during acoustic levitation, thus preventing compartmental mixing. Secondly, we demonstrate that the protein pore, alpha-hemolysin (aHL) and bacterial mechanosensitive ion channel of large-conductance (MscL), may be reconstituted into the membranes of levitated ACDC droplets to enable ionic mediated communication between selected compartments (Figure 4). The calcium sensitive fluorescent dye, Fluo-8, is incorporated within a droplet also containing monomeric alpha-hemolysin, which is connected to Ca^2+^ containing droplets within the internal core network (Fig.4 A-1). An additional Fluo-8 containing droplet without alpha-hemolysin is also connected to calcium containing droplets and encapsulated within the ACDC droplet, serving as a control. Fluo-8 fluorescence in the respective cores of the levitated ACDC droplet shows calcium flux through the membrane spanning aHL pores, whereas the non-aHL containing membrane remains impermeable (Fig.4 A-2). The observed sigmoidal fluorescent response is a consequence of an acceleration in ion-flux as an increasing number of functional heptameric pores assemble on the membrane from their constituent monomers [44]. The fluorescence response then plateaus (~20 mins) as the Fluo-8 becomes saturated (more discussion available in Fig. S7, Fig. S8).

**Fig 4.**
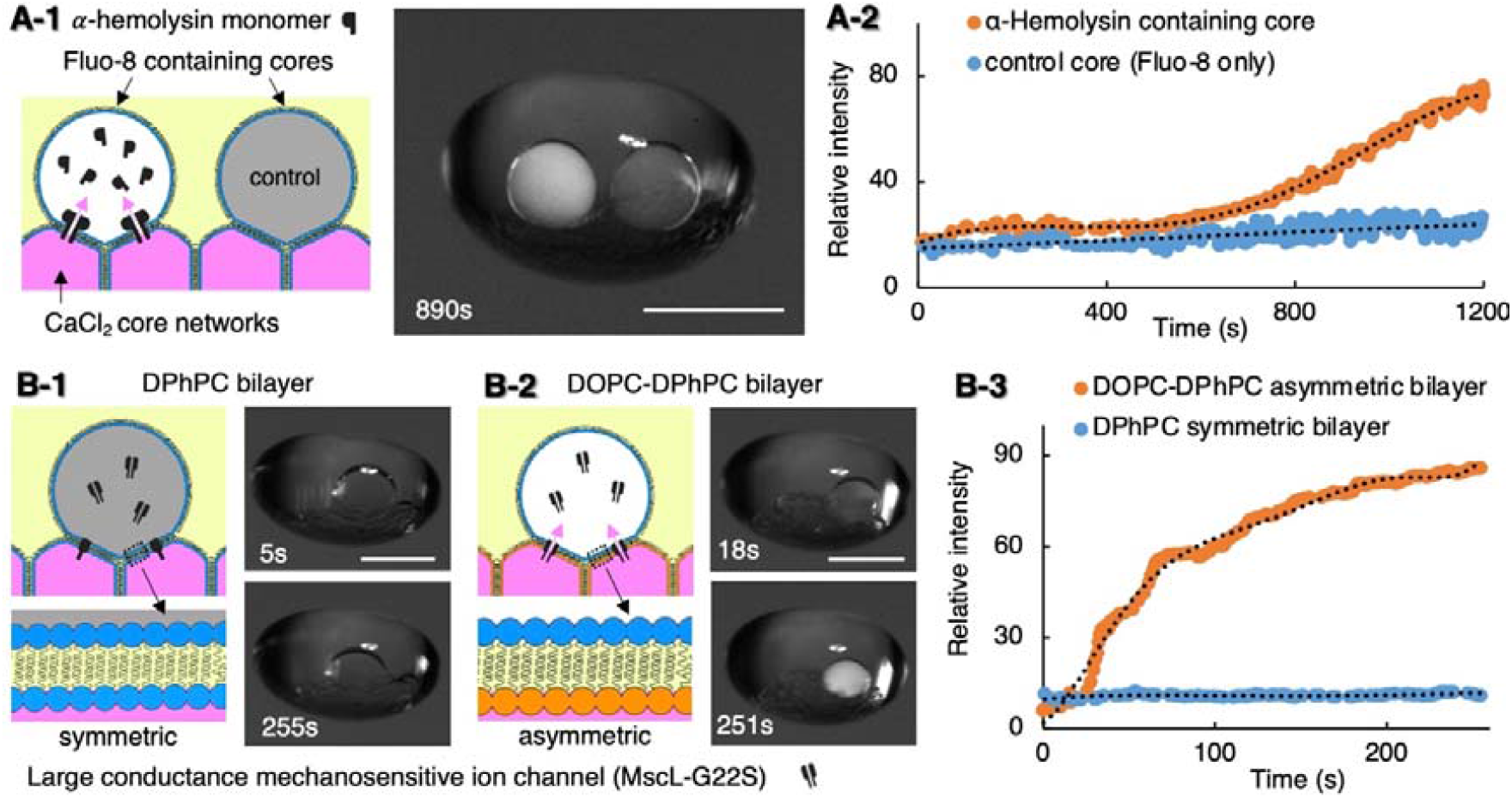
Reconstitution of functional alpha-hemolysin pores and bacterial MscL protein channels into the artificial membranes of the core network of levitated ACDC droplets. **A-1**. Alpha-hemolysin (aHL) monomers contained in an internal core bind to the membrane separating neighbouring droplets and assemble into heptameric protein pores in the DPhPC bilayer. Ca^2+^ ions diffuse through aHL pores moving from the source cores to the protein containing core where fluorescence is induced by the presence of the calcium sensitive fluorophore, Fluo-8. No ion-flux is observed in an identical control droplet without alpha-hemolysin monomer. **A-2**. The graph indicates the fluorescence intensity increase over time of both aHL containing and non-aHL containing (control) droplets in the levitated ACDC droplet over 20 minutes. (Dotted lines = moving average). **B**. The mechanosensitive channel, MscL, opens in response to membrane tension. Gating of MscL channels in levitated ACDC droplets with membranes of different bilayer leaflet compositions. **B-1**, In symmetric DPhPC bilayer networks no Ca^2+^ flux is observed, the channel remains closed; **B-2**, In asymmetric DOPC-DPhPC bilayer networks asymmetric membrane tension is induced, opening MscL channels, and giving rise to Ca^2+^ flux and observed fluorescence. (No fluorescence increase is observed in the absence of MscL (SI)) **B-3**, Time-course fluorescence in symmetric (DPhPC) and asymmetric (DOPC-DPhPC) membranes as depicted in B-1 and B-2. Asymmetric membrane tension induced activation of MscL channels enables Ca^2+^ flux, eliciting maximal fluorescence response over the course of 250 s. No fluorescence increase is observed in the symmetric membrane system indicative of channel inactivation. Scale bars = 1 mm.

Unlike the permanently open alpha-hemolysin pore that allows the diffusion of Ca^2+^ following pore formation, the mechanosensitive ion channel, MscL, is typically closed but can be gated by the application of mechanical stress applied to the membrane [45, 46]. A glycine-serine substitution (G22S) in the hydrophobic channel gate generates a mutant that is not spontaneously active, but possess a lower activation threshold than wild-type MscL [47, 48] Interestingly, on incorporation into ACDC droplets with membranes formed of DPhPC lipid (as used for aHL), no ion-flux is observed, indicating that the MscL channels are not spontaneously activated by the acoustic levitation or any associated acoustic perturbation of the membrane network (Fig 4B-1). The formation of asymmetric DPhPC-DOPC membranes between contacting cores of the ACDC droplet could be harnessed to induce membrane tension and consequently spontaneous MscL channel activation (Fig 4B-2). In these asymmetric membranes, rapid ion flux was observed via Fluo-8 fluorescence, reaching maximal response within 4 minutes (Fig. 4B-3). No measurable ion-flux was detected in the DPhPC symmetric membrane system. Asymmetric DPhPC-DOPC lipid bilayers were constructed by the *in situ* addition of a DPhPC monolayer coated, MscL containing, core into the levitated ACDC droplet with a DOPC-based core network. The DPhPC MscL containing core was initially prepared by incubating a MscL containing aqueous droplet in a DPhPC containing oil phase, to form a DPhPC monolayer coated core. Contact between the DOPC core network and the manually added DPhPC core allowed for the formation of an asymmetric bilayer with one leaflet mainly consisting of DPhPC and the other mainly consisting of DOPC. Once the DPhPC-DOPC bilayer is formed, the MscL channels became reconstituted and passively activated in the asymmetric membranes. Similar results were also achieved in the levitated ACDC droplet by the inclusion of LysoPC within the MscL containing core of a symmetric DPhPC bilayer membrane, with LysoPC being incorporated into the lipid bilayer to induce membrane tension and protein activation [46].

### Remote control of protein function in levitated ACDC droplet to activate intracellular communication

The ability to modulate protein channel function, on demand, represents an attractive proposition for the attainment of greater functional control over artificial cells. In this context, we demonstrate that the gating of MscL channels can be selectively activated on demand via remote perturbation using a combination of applied acoustic and magnetic forces (Fig. 5, SI Video 3). In symmetric DPhPC membranes within an ACDC droplet, the MscL channel is closed (Fig. 4B-1). The inclusion of a dedicated core within the droplet network containing magnetic particles, enables the orientation and ‘locking’ in position of the ACDC droplet on application of an external magnetic field to the levitated droplet (Fig. 3C).

**Fig 5.**
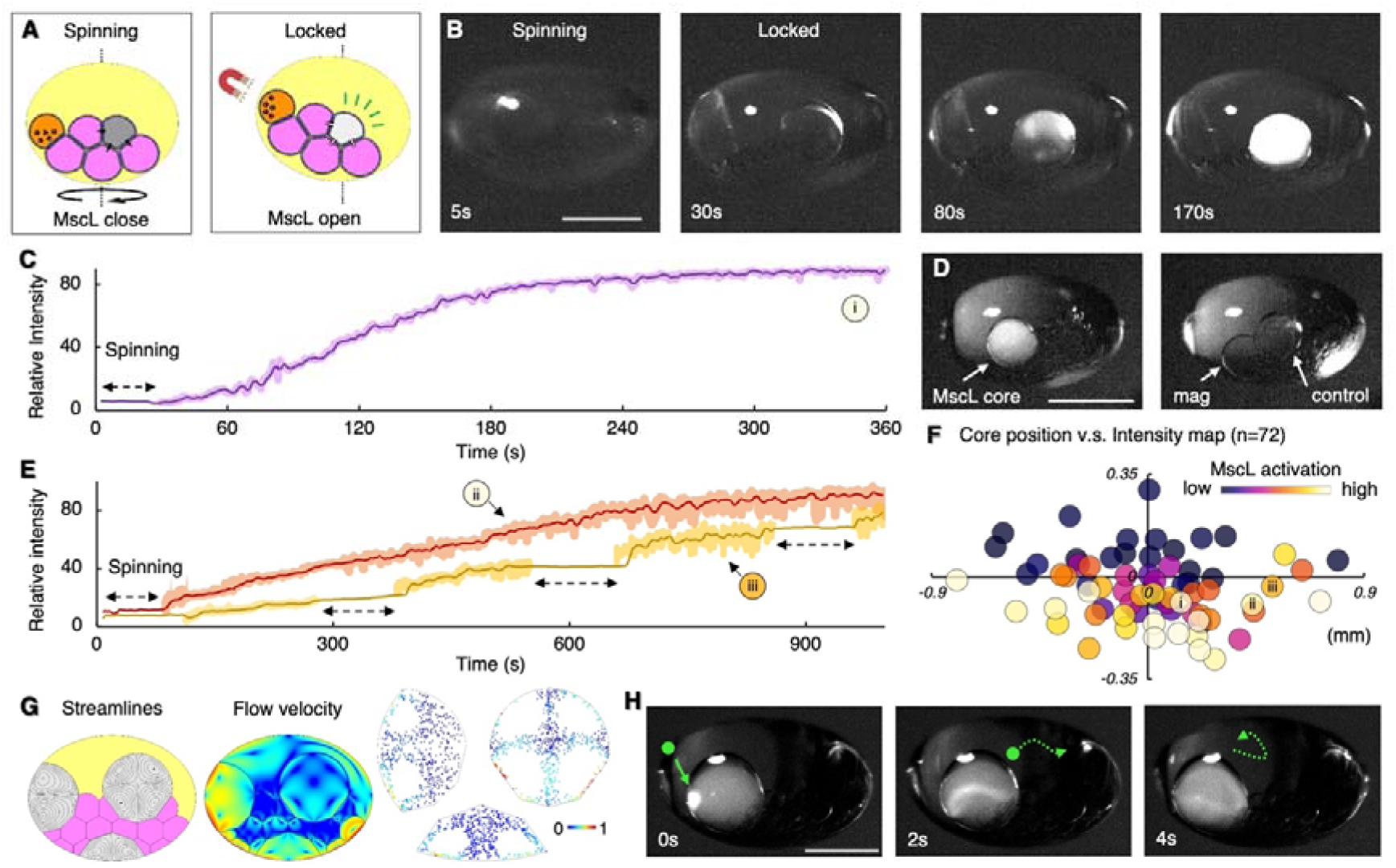
Remote control of ion channel gating in levitated ACDC droplet core networks using magnetic manipulation. **A**. Experimental concept: The incorporation of a core containing magnetic particles enables switching of the levitated ACDC capsule from a freely rotating to a locked state on the application of an external magnetic field. In the free (spinning) state acoustic forces are dissipated by subtle movement. In the locked state, this is dampened and instead manifests as induction of membrane tension. This may be used to selectively activate MscL channel gating. **B**. Time sequence images of MscL gating in a levitated ACDC droplet. From left to right; t = 5 s in the absence of an applied magnetic field the ACDC droplet is allowed to spin in the acoustic trap. At t = 30 s the magnetic field is applied, locking the ACDC droplet in position. MscL is activated and Ca^2+^ ion flux induced, observed by the fluorescent reporter, fluo-8, present in the MscL containing core. At t= 80 s, significant fluorescence increase is observed. With the fluorescence response saturated at t = 170s. In the absence of MscL no fluorescence increase is observed (Fig. S9). **C**. Fluorescence intensity trace of Ca^2+^ flux through MscL channels into a fluo-8 containing core of a locked, levitated, ACDC droplet. (Same experiment labelled (i) in Fig. 5e). (solid line = moving average). **D**. Images of a levitated ACDC droplet with a MscL containing core (left), a magnetic particle containing core and a control core containing Fluo-8 only (right). The orientation of the droplet here is controlled by the placement of external magnet to enable image acquisition from two angles to illustrate the three highlighted cores. **E**. Repeat activation/deactivation of MscL gating in levitated ACDC droplets can be achieved by repeated lock/spin operations. The orange trace shows an ACDC droplet experiencing one cycle of spin/lock operation, activating the ACDC capsule and enabling ionic communication. (Same experiment labelled (ii) in Fig. 5e). The yellow trace shows the ACDC droplet experiencing four sequential cycles of spin/lock operation, enabling repeated activation and shut-down of ionic communication within the ACDC capsule. Each spin interval was controlled as 100 s in duration, and the relative fluorescence intensity did not increase during the spin interval, indicating the MscL channels were mostly deactivated during these periods and activated on application of the magnetic lock. (Same experiments labelled (iii) in Fig. 5e). **F**. Relationship of core position and MscL activity: Heatmap depicting the relative rate of ion flux (rate of fluorescence increase) for MscL protein containing cores (colour) and their relative positions within different ACDC droplets (x/y offset from ACDC capsule centre). The plot origin is the centroid of each levitated ACDC droplet, each circular data point represents a separate experiment with the circle’s coordinates corresponding to the position of each protein containing core relative to the center of the ACDC droplet it is contained within. Marker colour indicates relative activity. The extent of channel activation is observed to correlate with position, with droplet membranes in the lower half and nearer the perimeter of the ACDC capsule displaying greater activity. Markers i, ii and iii correspond to the individual traces shown in Fig. 5 c&e. **G**. Finite element modelling of acoustic streaming induced convective flows in three simulated protein containing cores (grey) in different positions of a locked levitated ACDC droplet. Left: acoustically induced convective flow streamlines in the protein cores located at three different positions. Middle: flow velocity magnitude (logarithmic scale) within the cores and oil phase of the simulated ACDC droplet. Right: Simulated distribution of 5 μm particles due the flow within the protein containing cores. These simulations show how acoustic streaming pressure varies with core position, corroborating experimental findings in Fig 5F **H**. Experimental time sequence images over 4 s show local Ca^2+^ release through MscL channels at the membrane followed by dissipation of the fluorescence following the acoustic streaming flow (corresponding to flow simulations in Fig 5G). Such hot-spots of MscL activation were observed stochastically in regions and periods of high induced local tension. Green spots and arrow dotted lines indicate the flow direction. Scale bars = 1 mm.

In a non-locked state (Fig. 4B-1), no ion flux is observed as the MscL channels remain in a closed state. With the incorporation of the magnetic particles in the core, in the absence of a magnetic field, the MscL channels remain closed (0-30 s Fig 5B & C). The MscL channels are only activated on application of an external magnetic field to ‘lock’ the droplet, with ion flux only observed following the application of the external magnetic field (30-240s Fig 5B & C). The fluorescent response rapidly proceeds to maximum within ~3 minutes of activation, indicating a comparable extent of channel opening to that observed with formation of a DPhPC-DOPC asymmetric membrane. In the absence of MscL, no fluorescence intensity increase is observed in the locked, Fluo-8 containing, core (Fig 5D), evidencing the membranes remaining intact with the ion-flux being protein mediated (Fig. S9). By the selective application of the external magnetic field, to move between a ‘locked’ (MscL active) and unlocked/rotating (MscL inactive) states, it is possible to repeatedly control MscL opening and closing in the ACDC artificial cell (Fig. 5E). Unlike chemical activation, this approach enables deactivation and repeat activation, creating opportunities for multiple use switches analogous to that of biological cells, rather than one-time only activation typically employed in artificial cell communication and processes initiated on the addition or liberation of activating species. This selective on/off activation is possible due to the inability of the magnetically ‘locked’ droplet system to effectively redistribute the acoustic irradiation stress as the total force remains, which would otherwise be minimised through small changes in orientation and position. Acoustic irradiation stress has previously been applied to deform lipid vesicles and small living organisms to measure the membrane Young’s modulus [49], whilst boundary-induced acoustic streaming effects were studied, indicating steady-state torque can be induced on levitated samples by applying asymmetry in the system [38]. In our experiment, without magnetic ‘locking’, the asymmetry of the ACDC droplet architecture, levitator configuration, and other environmental and operational forces exerted on the ACDC droplet, manifest asymmetrically and collectively to induce the torque which can be redistributed by subtle movement of the levitated ACDC droplet. This is due to the low friction within the system, and usually observed as re-centring (Fig. 1 B & C), convective flows (Fig. 1D), and slow rotation (Fig. 3C). In the ‘locked’ configuration, such energy cannot covert to kinetic motion but manifests as the membrane tension within the core of the network, sufficient to activate the incorporated MscL channels (Fig. 5).

The activation of the MscL channel was found to be influenced by the protein containing core position within the levitated ACDC droplet (Fig 5F). MscL does not tend to activate when the protein containing core is close to the top or within the mid half of the ACDC droplet, whereas greater MscL activation is achieved when the protein containing core is near the bottom half or near the perimeter of the ACDC droplet. We rationalise that this is likely due to the higher magnitudes of convective flow velocity within the cores in these locations, as shown by numerical modelling (Fig 5G, SI Video 3), resulting in a greater shear rate of the fluid flow, and hence greater tension applied to the lipid bilayers located in these regions. In addition, as shown in Fig 5H, rapid and large Ca^2+^ fluxes originated at local membranes as an indication of mass MscL gating, with the fluorescence then dissipating through the protein containing core following the convective flow induced by the acoustic streaming effect (Fig 5H). These flows are compatible with simulated convection and tracer microparticle distributions (Fig 5G). We again rationalise that the stochastic nature of these mass-activation events is, in part, due to the interplay of precise membrane position, local variations in acoustic pressure and the ability of the wider core network to redistribute induced tension, creating transient local ‘hot-spots’ of high channel activation. This phenomenon was not observed in the aHL group, where ion-flux is a consequence of the more homogeneous passive diffusion of ions through the formed aHL pores, which is not modulated in this way.

In principle, this MscL gating behaviour could be used to control the generation of chemical gradients, whilst the internal acoustic streaming may govern the dynamics of mixing of encapsulated contents in specific cores for selected chemical reactions, both enabling more precise core network programming and control. In the MscL activation experiments, whilst the rate of fluorescence intensity increase depends on the membrane position and application of the dual magnetic and acoustic perturbation, it is also influenced by the ionic gradient and the exact reconstituted protein densities, which is itself a function of bilayer area, droplet volume, protein concentration and reconstitution efficiency [50]. Protein reconstitution efficiency can be a source of variability, but with control over these variables alongside core network architecture, it indicates that more sophisticated control of ion signalling via controlled activation/deactivation of MscL can be achieved. Overall, functional membrane proteins can be reconstituted in the lipid bilayer membranes of levitated ACDC droplets, with aHL and MscL used here to dictate ionic transfer within the core networks of a levitated ACDC droplet. MscL channels can be remotely activated/deactivated, responding to a contactless magnetic operation in an acoustic standing wave, for on-demand and repeatable remote operation of chemical signalling in artificial cells.

## Discussion

In summary, we present a new platform for the creation, manipulation, control and measurement of complex-emulsion-based artificial cells (ACDC droplets) at single compartment (core) resolution, using droplet microfluidics and acoustic levitation. The sizes, reagent encapsulations, and 3D organisations of cores can be controlled by programmed microfluidic circuits and inlet flow profiles, with droplet sequence determining 3D droplet network connectivity. The levitated artificial cell droplets can be operated in situ to reconfigure the compartmentalised droplet network in the acoustic levitator, orientate the structure on-demand, and apply thermal and light energies. This is demonstrated by pneumatic spinning, magnetic manipulation, heat irradiation and free-radical photopolymerization, respectively. We show that lipid bilayers segregating the internal compartments of the artificial cell, are robust and leak-free in the acoustic field. Furthermore, we demonstrate the ability to reconstitute alpha-hemolysin and MscL membrane proteins, to direct Ca^2+^ ion transport across membranes segregating select cores of the levitated compartmentalised droplet. Importantly, the gating of incorporated MscL channels can be selectively and repeatedly remote controlled to activate/deactivate channel activity through the application of an external magnetic field alongside the levitating acoustic field. In this way membrane tension can be perturbed to induce channel opening. This establishes remotely-controlled intracellular communication within the artificial cell, which can be repeatedly switched on and off. Such a combination of acoustic and magnetic field activation could be used to activate artificial cells *in situ* for future diagnostic and therapeutic applications, or used to initiate and halt artificial cell functions. The components of the presented experimental platform are all fabricated using accessible 3D-printing and have robust performance in diverse settings, hence this approach is both generalisable and reconfigurable for different chemical and synthetic biology laboratories.

Structuring intra-cellular compartments of artificial cells with organised chemistries is an important milestone in the bottom-up creation of artificial cells affording increased functionality. The integration of droplet forming, sequencing, and encapsulating microfluidics alongside post processing, affords a greater control space for the fabrication and operation of individual complex emulsion droplets as artificial cells. Following *in situ* processing, such droplets can be subsequently 3D-printed to construct more complicated, tissue-like superstructure materials. This work provides a versatile platform technology for engineering multicompartment artificial cell chassis, that bridge biomimetic materials and cell functionality development. With the developed platform, it is possible to investigate and operate artificial cells at varying length scales, ranging from the building blocks, such as proteins and lipid bilayers, to the droplet network organisation and architecture, whilst the function and morphology can be altered, or activated, repeatedly providing analogous processes to those of natural cells, in comparison to typical approaches with one-time activation and subsequent attainment of equilibrium. The ability to activate and deactivate membrane proteins to govern reaction or chemical communication pathways on demand provides a pathway to the more routine use of artificial cells as next generation smart materials. In this context, the demonstrated capability will enable the design of programmable cell behaviours, incorporating both multiphysics engineering and (bio)chemical approaches, for example molecular assembly and biochemical synthesis [51, 52, 53] could be incorporated and integrated with the programmable reconfiguration and remote activation demonstrated. Such advances will enhance our capability to create and control increasingly complex artificial cells and potential artificial biological systems [54,55,56], comprised of functional organelle-like sub-compartments in active chemical communication, towards the development of more sophisticated hierarchical, life-like materials and soft matter smart materials.

## Methods

All the experiment and methods details are available in Supplementary Information.

## Author Contributions

J.L. conceived and led the research, and performed all the experiment. O.K.C., B.W.D., and D.A.B equally contributed to the experiment plan designed by J.L.. D.A.B. contributed to the provision of microfluidic manifold structures. W.D.J. and O.K.C. contributed to protein assay and fluorescence imaging. O.K.C. conceived the ion channel activation experiments and developed these and their analysis together with W.D.J. and J.L. O.K.C designed and built the optical imaging setup with W.D.J. B.W.D. built the acoustic levitator and contributed to levitator experiments. P.D., B.W.D. and D.A.B. contributed to Multiphysics simulation. W.X contributed to data analysis. P.R., M.B. and B.M. produced and purified the MscL-G22S as the key reagents. J.L., O.K.C., and D.A.B. wrote the manuscript. All authors discussed the results and commented on the manuscript. The authors declare that they have no competing interests.

## Acknowledgements

This work was primarily supported by funded by the European Horizon 2020 project ACDC (Artificial Cells with Distributed Cores) under project award number 824060 awarded to D.A.B and O.K.C at Cardiff University. The custom-built fluorescent imaging setup and development was in part supported by a Cardiff University MRC Confidence in Concept award (MC_PC_17170) awarded to O.K.C. B.W.D. would like to acknowledge financial support from the Wolfson Foundation and the Royal Society.

## Additional information

Supplementary Information is available for this paper online.

## References

1. Martin, W. et al. Evolutionary origins of metabolic compartmentalization in eukaryotes. Philosophical Transactions of the Royal Society B: Biological Sciences 365, 847–855 (2010).

2. Ball, P. Synthetic biology: starting from scratch. Nature 431, 624–627 (2004).

3. Schwille, P., et al. MaxSynBio: avenues towards creating cells from the bottom up. Angewandte Chemie International Edition 57, 13382–13392 (2018).

4. Deshpande, S., & Dekker, C. On-chip microfluidic production of cell-sized liposomes. Nature protocols 13, 856–874 (2018).

5. Göpfrich, K., et al. One-pot assembly of complex giant unilamellar vesicle-based synthetic cells. ACS synthetic biology 8, 937–947 (2019).

6. Leptihn, S., et al. Constructing droplet interface bilayers from the contact of aqueous droplets in oil. Nature protocols 8, 1048–1057 (2013).

7. Sakuta, H., et al. Self-Emergent Protocells Generated in an Aqueous Solution with Binary Macromolecules through Liquid-Liquid Phase Separation. ChemBioChem 21, 3323 (2020).

8. Vance, J. A., & Devaraj, N. K. Membrane Mimetic Chemistry in Artificial Cells. Journal of the American Chemical Society (2021).

9. Niederholtmeyer, H., Chaggan, C., & Devaraj, N. K. Communication and quorum sensing in non-living mimics of eukaryotic cells. Nature communications 9, 1–8 (2018).

10. Joesaar, A., et al. DNA-based communication in populations of synthetic protocells. Nature nanotechnology 14, 369–378 (2019).

11. Buddingh, B. C., Elzinga, J., & van Hest, J. C. Intercellular communication between artificial cells by allosteric amplification of a molecular signal. Nature communications 11, 1–10 (2020).

12. Tayar, A. M., Karzbrun, E., Noireaux, V., & Bar-Ziv, R. H. Synchrony and pattern formation of coupled genetic oscillators on a chip of artificial cells. Proceedings of the National Academy of Sciences 114, 11609–11614 (2017).

13. Elani, Y. Interfacing living and synthetic cells as an emerging frontier in synthetic biology. Angewandte Chemie International Edition, 60, 5602–5611 (2021).

14. Mukwaya, V., et al. Programmable Membrane-Mediated Attachment of Synthetic Virus-like Nanoparticles on Artificial Protocells for Enhanced Immunogenicity. Cell Reports Physical Science 2, 100291 (2021).

15. Zhou, L., et al. Lipid-bilayer-supported 3D printing of human cerebral cortex cells reveals developmental interactions. Advanced Materials 32, 2002183 (2020).

16. Liu, S., et al. Enzyme-mediated nitric oxide production in vasoactive erythrocyte membrane-enclosed coacervate protocells. Nature Chemistry 12, 1165–1173 (2020).

17. Martin, N., Douliez, J. P., Qiao, Y., Booth, R., Li, M., & Mann, S. (2018). Antagonistic chemical coupling in self-reconfigurable host–guest protocells. Nature communications, 9(1), 1–12.

18. Mu, W. et al. Membrane-confined liquid-liquid phase separation toward artificial organelles. Science advances 7, eabf9000 (2021).

19. Aufinger, L., & Simmel, F. C. (2018). Artificial gel-based organelles for spatial organization of cell-free gene expression reactions. Angewandte Chemie International Edition, 57(52), 17245–17248.

20. Cazimoglu, I., Booth, M. J., & Bayley, H. A lipid-based parallel processor for chemical signals. (Preprint at https://www.biorxiv.org/content/10.1101/2021.05.05.442835v1.full.pdf+html) (2021).

21. Einfalt, T., et al. Bioinspired molecular factories with architecture and in vivo functionalities as cell mimics. Advanced Science 7, 1901923 (2020).

22. Xu, D., Kleineberg, C., Vidaković-Koch, T., & Wegner, S. V. Multistimuli Sensing Adhesion Unit for the Self-Positioning of Minimal Synthetic Cells. Small 16, 2002440 (2020).

23. Booth, M. J., Cazimoglu, I., & Bayley, H. Controlled deprotection and release of a small molecule from a compartmented synthetic tissue module. Communications Chemistry 2, 1–8 (2019).

24. Miller, T. E., et al. Light-powered CO2 fixation in a chloroplast mimic with natural and synthetic parts. Science 368, 649–654 (2020).

25. Alcinesio, A. et al. Controlled packing and single-droplet resolution of 3D-printed functional synthetic tissues. Nature communications 11, 1–13, (2020).

26. Booth, M. J., Schild, V. R., Graham, A. D., Olof, S. N., & Bayley, H. Light-activated communication in synthetic tissues. Science Advances 2, e1600056 (2016).

27. Downs, F. G., et al. Multi-responsive hydrogel structures from patterned droplet networks. Nature chemistry 12, 363–371 (2020).

28. Elani, Y., Gee, A., Law, R. V., & Ces, O. Engineering multi-compartment vesicle networks. Chemical Science 4, 3332–3338 (2013).

29. Villar, G., Graham, A. D., & Bayley, H. A tissue-like printed material. Science 340, 48–52 (2013).

30. Villar, G., Heron, A. J., & Bayley, H. Formation of droplet networks that function in aqueous environments. Nature nanotechnology 6, 803–808 (2011).

31. Kwok, R. Five hard truths for synthetic biology. Nature News 463, 288–290 (2010).

32. Decoene, T. et al. Standardization in synthetic biology: an engineering discipline coming of age. Critical reviews in biotechnology, 38, 647–656 (2018).

33. Deng, N. N., & Huck, W. T. Microfluidic formation of monodisperse coacervate organelles in liposomes. Angewandte Chemie 129, 9868–9872 (2017).

34. Linsenmeier, M. et al. Dynamics of synthetic membraneless organelles in microfluidic droplets. Angewandte Chemie 131, 14631–14636 (2019).

35. Deng, N. N., Yelleswarapu, M., Zheng, L., & Huck, W. T. Microfluidic assembly of monodisperse vesosomes as artificial cell models. Journal of the American Chemical Society 139, 587–590 (2017).

36. Marzo, A., Barnes, A., & Drinkwater, B. W. TinyLev: A multi-emitter single-axis acoustic levitator. Review of Scientific Instruments 88, 085105 (2017).

37. Tian, L., Li, M., Patil, A. J., Drinkwater, B. W., & Mann, S. Artificial morphogen-mediated differentiation in synthetic protocells. Nature communications 10, 1–13 (2019).

38. Trinh, E. H., & Robey, J. L. Experimental study of streaming flows associated with ultrasonic levitators. Physics of Fluids 6, 3567–3579 (1994).

39. Baxani, D. K. et al. Bilayer networks within a hydrogel shell: a robust chassis for artificial cells and a platform for membrane studies. Angewandte Chemie International Edition 55, 14240–14245 (2016).

40. Li, J. et al. Formation of Polarized, Functional Artificial Cells from Compartmentalized Droplet Networks and Nanomaterials, Using One-Step, Dual-Material 3D-Printed Microfluidics. Advanced Science 7, 1901719 (2020).

41. Cho, E., & Lu, Y. Compartmentalizing Cell-Free Systems: Toward Creating Life-Like Artificial Cells and Beyond. ACS Synthetic Biology 9, 2881–2901 (2020).

42. Müller, A., Schneider, J. J., & Schömer, E. Packing a multidisperse system of hard disks in a circular environment. Physical Review E 79, 021102 (2009).

43. Lim, M. X., Souslov, A., Vitelli, V., & Jaeger, H. M. Cluster formation by acoustic forces and active fluctuations in levitated granular matter. Nature Physics 15, 460–464 (2019).

44. Thompson, J. R., Cronin, B., Bayley, H., & Wallace, M. I. Rapid assembly of a multimeric membrane protein pore. Biophysical journal 101, 2679–2683 (2011).

45. Perozo, E., Cortes, D. M., Sompornpisut, P., Kloda, A., & Martinac, B. Open channel structure of MscL and the gating mechanism of mechanosensitive channels. Nature 418, 942–948 (2002).

46. Strutt, R., et al. Activating mechanosensitive channels embedded in droplet interface bilayers using membrane asymmetry. Chemical Science 12, 2138–2145 (2021).

47. Rosholm, K. R. et al. Activation of the mechanosensitive ion channel MscL by mechanical stimulation of supported Droplet-Hydrogel bilayers. Scientific reports, 7, 1–10, (2017).

48. Yoshimura, K., Batiza, A., Schroeder, M., Blount, P., & Kung, C. Hydrophilicity of a single residue within MscL correlates with increased channel mechanosensitivity. Biophysical journal 77, 1960–1972 (1999).

49. Silva, G. T. et al. Acoustic deformation for the extraction of mechanical properties of lipid vesicle populations. Physical Review E 99, 063002 (2019).

50. Castell, O. K., Berridge, J., & Wallace, M. I. Quantification of membrane protein inhibition by optical ion flux in a droplet interface bilayer array. Angewandte Chemie International Edition 51, 3134–3138 (2012).

51. Hua, Z., et al. Anisotropic polymer nanoparticles with controlled dimensions from the morphological transformation of isotropic seeds. Nature communications 10, 1–8 (2019).

52. Nakatani, N. et al. Specific Spatial Localization of Actin and DNA in a Water/Water Microdroplet: Self-Emergence of a Cell-Like Structure. ChemBioChem 19, 1370 (2018).

53. Song, Y. et al. Budding-like division of all-aqueous emulsion droplets modulated by networks of protein nanofibrils. Nature communications 9, 1–7 (2018).

54. Beneyton, T. et al. Out-of-equilibrium microcompartments for the bottom-up integration of metabolic functions. Nature communications 9, 1–10 (2018).

55. Li, Q., Li, S., Zhang, X., Xu, W., & Han, X. Programmed magnetic manipulation of vesicles into spatially coded prototissue architectures arrays. Nature communications 11, 1–9 (2020).

56. Samanta, A., Sabatino, V., Ward, T. R., & Walther, A. Functional and morphological adaptation in DNA protocells via signal processing prompted by artificial metalloenzymes. Nature nanotechnology 15, 914–921 (2020).

