## supplementary material for "Building programmable multicompartment artificial cells incorporating remotely activated protein channels using microfluidics and acoustic levitation"

**Content of supplementary materials**

**Experimental Methods**

**Fig S1. Droplet laboratory experiment setup**

**Fig S2. Experimental and simulated droplet trajectories in the acoustic trap**

**Fig S3. The relationship between levitator input voltage and the levitated droplet shape**

**Fig S4. Simulated and experimental details of 3D-printed, multi-layered, droplet forming fluidic junctions.**

**Fig S5. Complementary data of pneumatic operations for the spinning of levitated ACDC droplets (Fig 3B).**

**Fig S6. Shell photo-polymerisation of a microfluidically-formed, levitated oil/PEGDA multi-phase droplet.**

**Fig S7. Relative fluorescent intensity of the Fluo-8 containing cores (white) with different concentrations of pre-loaded CaCl_2_ in levitated ACDC droplets.**

**Fig S8. Evaporation effects on fluorescence intensity are minimal compared to Ca^2+^ transport over the course of 30 minutes.**

**Fig S9. Relative intensity of a control core (white dotted circle) containing fluo-8 but no MscL, in a levitated ACDC droplet.**

**SI Video 1. Levitation of ACDC droplet**

**SI Video 2. Contactless operation of ACDC droplet**

**SI Video 3. MscL protein channel activation**

**Experimental Methods**

**Chemicals and Components:** Alginic acid sodium salt, calcium chloride, potassium chloride, silicone oil AR20, hexadecane, glucose, HEPES, EDTA, PEGDA, sulforhodamine B, calcein, lissamine green, alpha-hemolysin (aHL) monomer and n-dodecyl β-D-maltoside (DDM) were all purchased from Sigma-Aldrich. Fluo-8H AM was purchased from ATT Bioquest. BioRad Chelex 100® resin was purchased from BioRad. Hydrophilic, magnetic silica microparticles (SiMAG/MP-DNA 1.0 μm) were purchased from Chemicell. 1,2-Dioleoyl-sn-glycero-3-phosphocholine (DOPC) and 1,2-Diphytanoyl-sn-glycero-3-phosphocholine (DPhPC) lipids were purchased from Avanti Polar Lipids. Neodymium magnets were purchased from RS Components.

The preparation of the precursors for the ACDC droplets was as follows. Water phase: potassium chloride was dissolved in deionised water at 0.15 M concentration with 0.01 M HEPES and 150 μM water soluble dyes (sulforhodamine B, calcein, or lissamine green). Buffer was filtered with Nylon membrane filters (0.22 μm, Fisherbrand) for use in the experiment and stored as a stock solution. Oil phase: Lipid powders were dissolved in hexadecane at 20 mg mL^-1^ as stock, and mixed with additional hexadecane and silicone oil at different ratios for microfluidics experiments. To prepare the microparticle containing aqueous phase solution, the original particle-containing solution (from supplier at 50 mg mL^-1^) was mixed in the prepared 0.15 M KCl buffer at a ratio of 1:100 v/v. To prepare alginate solution, alginate powder was added to 0.15 M KCl buffer at 2% w/v, and mixed using a magnetic stirrer (IKA RCT basic safety control) agitated at 800 rpm, and at 50 °C, for 4 hr. All prepared solutions were kept for a maximum of one week for use in experiments.

**MscL-G22S expression and purification:** The MscL-G22S mutant expression construct was made from the wild-type MscL 3.1 construct using QuikChange site-directed mutagenesis kit (Agilent Technologies, Santa Clara, CA, USA) [1]. To purify the channel protein, MscL-G22S was expressed in BL-21 (DE3) (Novagen) E. coli strain, which was grown at 37 °C to OD600 0.8. The protein expression was induced with 1 mM IPTG for 3 h as previously described (Petrov et al., 2011). Briefly, the cell pellet was suspended in PBS in the presence of ~0.02 mg/mL DNase (Sigma DN25) and 0.02% PMSF (Amresco M145) and disrupted using a TS5/48/AE/6 A cell disrupter (Constant Systems) at 31,000 psi at 4 °C. Cell debris was removed by centrifugation at 12,000 × g for 15 min at 4 °C. The membranes were pelleted at 45,000 RPM in a 45 Ti rotor (Beckman) for 3 h at 4 °C. The pellets were then solubilized in PBS with 8 mM DDM overnight at 4 °C followed next day by centrifugation at 12,000 × g for 20 min at 4 °C. The supernatant containing the solubilized MscL G22S protein was incubated with cobalt sepharose (Talon®, 635502, Clontech) followed by several washes with PBS supplement containing 15 mM Imidazole (Sigma, 56750). The MscL protein was then eluted with 500 mM imidazole PBS and concentrated using a 100 kDa Amicon-15 centrifugal filter unit (Merck Millipore) by diluting imidazole with DDM PBS prior to centrifugation. Protein concentration was estimated using polyacrylamide electrophoresis with SimplyBlue™ (LC6065, Thermo Fisher) staining.

**Preparation of aqueous droplet solutions for protein related experiment:** Calcium chloride solutions used in the protein mediated communication experiments consisted of 0.89M CaCl_2_ and 10mM HEPES, adjusted to pH7.4. Protein containing and non-protein containing control droplets were made up from a base solution consisting of KCl, sucrose, EDTA and HEPES adjusted to pH7.4. BioRad Chelex 100 resin at ~5% w/v ratio was added to the base solution to remove any divalent cations which might affect the background fluorescence values on addition of Fluo-8H. The mixture was agitated on a tube roller for 2 hours before being filtered through a 0.2µM filter to remove the Chelex resin. To this Fluo-8H was added either alone, or along with either aHL, MscL1-G22S, or DDM (the detergent used to stabilise purified MscL) in the case of control droplets. The solution was made up to a final concentration of 1M KCl, 0.6M sucrose, 5µM EDTA, 10mM HEPES, 55.8µM Fluo-8H and either 155nM aHL monomer (alpha hemolysin droplets), 4 µM DDM (DDM only control droplets) or a 1:1000 dilution of MscL1-G22S stabilised with 1mM DDM (providing a final DDM concentration in experimental droplets of 1 µM) (mechanosensitive droplets). The KCl/sucrose concentrations used ensured that protein/control droplets were osmotically, but not ionically, matched to the CaCl_2_ containing droplets.

**Droplet laboratory setup:** Microfluidic devices were designed and modelled with COMSOL Multiphysics (the dimensions of tubular channels are in the range of 120–500 μm diameter), and were printed using fused filament fabrication printers (Ultimaker 5) with cyclic olefin copolymer filament (Creamelt), using 0.25 mm AA print-cores. The print g-codes were programmed using Cura (version 4.8.0) software with customized settings. The layer height was controlled at 0.06 mm, and the printing speed was tuned at 20 mm s^−1^ with default printing temperatures (255 °C). Ultimaker material station and air filtering system were used to maximum the printing quality. Droplet content precursors were loaded in syringes (0.5, 1, 2.5, 5 mL, gas-tight, SGE Analytical Science), and were delivered to the microfluidic devices at a constant flowrate, through PEEK and PFA interconnects and FEP tubing, using syringe displacement pumps (KD Scientific, model 789200L). Precursors were also loaded in 15mL Falcon tubes, and were delivered to the microfluidic devices using controllable pressure profiles, programmed by wave functions using ELVEFLOW pressure pumps (OB1 MK3+).

For the levitation of ACDC droplets, a multi-emitter, single-axis acoustic levitator was used, with an array of Murata MA40S4S transducers of 40 kHz frequency, which generates acoustic waves in air at a wavelength of 8.65 mm. The electric signal originates from a driver board circuit with an Arduino NaNo (to generate square signals) and an L297N amplifier. More information on the electronics can be found in the original TinyLev article [3]. The TinyLev device is composed of two oppositely facing arrays of 36 transducers each (72 in total) and are arranged in 3 rings of 6, 12 and 18 transducers, forming a hexagonal pattern. The distance between the top and bottom arrays of transducers (trapping region) was 11cm. The transducers are fixed on a 3D-printed skeleton of the TinyLev. Once this device is assembled, the transducers can transform the electric signal received from the driving board circuit into acoustic power, which causes trapping of the ACDC droplets.

The fluidics, acoustics, optics and other operational elements were fixed on an optical board (30cm*30cm) with 3D-printed stands and holders, as the prototype of the droplet laboratory. The linear movement of the parts were manually controlled by micromanipulators and lab-jacks. The pneumatic device was composed of a 3D printed stand to fix 30 cm of plastic tubing (inner diameter 0.5 mm) and the nozzle was placed 3 cm away from the levitated droplets. Air was loaded in a 60mL plastic syringe connected to the tubing, and the air inflow rate was controlled by a syringe pump. The magnetic manipulation device was composed of a 3D printed stand to mount a metal bar, to which a 1 cm^3^ cubic neodymium magnet was attached at the end. The heating element was composed of customised copper coils, made from copper wire (1 mm diameter), attached to a soldering iron of which the temperature can be controlled between 100 °C to 400 °C. This was mounted on a metal pole. A UV torch (365nm wavelength) was used for the PEGDA photopolymerisation reactions.

The imaging setup for general experimental images and videos was composed of a horizontally placed Nikon SMZ745T microscope with a mounted high-speed camera (MegaSpeed, up to 1300 frame per second with 640*480 pixels), a USB digital camera with default tele tube placed at 45 degree, and 3 light sources, including one RGB LED lamp (RS), one white light LED bulb (Thorlabs), and one LED ring (default with USB camera). A thermal imaging camera (Micro-Epsilon) was used to capture the temperate profile. All the components were placed at fixed positions during the experiment. A Nikon AZ100 and a Nikon MM 800 were used to take the images of the droplets flowing in microfluidic device and the images of formed alginate microgels in a Petri dish filled with buffer, respectively.

**Fluorescence imaging:** A custom built optical set up was added to the Nikon SMZ745T stereomicroscope in order to carry out fluorescence imaging experiments. This consisted of an imaging arm with two F=100mm lenses. The first placed ~100 mm from the c-mount of the stereomicroscope served to collimate light exiting the stereomicroscope. This provided an infinity corrected region of the light path into which a bandpass filter centred at 525nm with a width of 50nm (Edmund optics) was placed. The second F=100mm lens was located after the filter ~100 mm from the exterior housing of a Basler Pulse pu1280-54um camera. Image capture was carried out using Basler’s pylon viewer software V6.2 with global shuttering on, an exposure time of 10ms and an acquisition rate of 25Hz. Illumination was achieved by selective use of blue LED array as part of the RGB LED light source (RS). Lenses, lens tubing and other optomechanic elements were purchased from Thorlabs.

**Data analysis:** Images and videos were analysed using Nikon Element software and the FIJI distribution [4] of ImageJ software with customised coding of Macros. Thermo profiles were analysed by TIM Connect software. Numerical data were processed with MATLAB (R2020a) and Microsoft Excel.

**Multiphysics simulation:** Computational models of the TinyLev acoustic field were achieved using COMSOL Multiphysics software (version 5.4). The structure of TinyLev (in the format of an STL file) was imported into COMSOL for the 3D simulation. The distribution of the acoustic field within the trapping region was solved using the Pressure Acoustics, Frequency Domain interface, while trapping of a 2mm aqueous droplet in air was modelled using the Particle Tracing for Fluid Flow Interfaces. The Pressure Acoustics, Frequency Domain Interface solves Helmholtz equations for given frequencies, detailed in the COMSOL Multiphysics acoustic module user’s guide, and within the frequency domain, the Helmholtz equation becomes Eq. (1):

$$-\frac{1}{\rho}\nabla^{2}p_{t}-\frac{k_{eq}^{2}}{\rho}p_{t}=0$$

(1)

$$p_{t}=p+p_{b}$$

$$k_{eq}^{2}=\left( \frac{\omega}{c} \right)^{2}$$

where $p_{t}$ is the total acoustic pressure, $k_{eq}$ is the wave number, $\rho$ is the density, and $c$ is the speed of sound. However, no Background Pressure Field was added to the model, hence $p_{t}=p_{0}$, where $p_{0}$ is a pressure variable defined within the interface.

When introducing the Particle Tracing for Fluid Flow interface for droplet trapping, it allows interaction between the acoustic field distribution from the first model (Pressure Acoustics, Frequency Domain) and particles introduced from the Particle Tracing interface.

For the acoustic streaming simulations, a 2D geometry was prepared, including models of the ACDC droplet domain and sub-compartments. Initially, the model uses the Thermoviscous Acoustics, Frequency Domain interface, which solves for the first order acoustic fields $\left( p_{1},u_{1},T_{1} \right)$. The boundaries of the ACDC droplet defined in this model will be responsible for the streaming effects. Next, the Laminar Flow Interface is added to solve the time-averaged net flow [5], driven by the first order acoustic fields as shown in Eq.(2) and Eq.(3):

$$\nabla\cdot\left( \rho_{0}\boldsymbol{u}_{\mathbf{2}} \right)=-\nabla\cdot\left( \left\langle\rho_{1}\boldsymbol{u}_{\mathbf{1}} \right\rangle\right)$$

(2)

$$\nabla\cdot\boldsymbol{\sigma}_{\mathbf{2}}=\nabla\cdot\left( \rho_{0}\left\langle\boldsymbol{u}_{\mathbf{1}}\boldsymbol{u}_{\mathbf{1}}^{\boldsymbol{T}} \right\rangle\right)$$

(3)

where the terms with subscript 1 belong to the first order acoustic field solution and terms with subscript 2 belong to the streaming flow model. The first order terms (R.H.S) are introduced into the Laminar Flow model as weak contributions in domains and boundaries. Τo complete the simulation of the streaming effect within the ACDC droplet, Stokes drift contributions are included on the boundaries responsible for the streaming effect, [5] in the form of Eq. (4):

|  | $u_{2}=-\left\langle\left( s_{1}\cdot\nabla\right)u_{1} \right\rangle$ | (4) |
| --- | --- | --- |
|  | where $s_{1}=\frac{u_{1}}{i\omega}$ |  |

Finally, particles of 5 µm diameter were released from certain cores using the Particle Tracing for Fluid Flow interface and their trajectories investigated under the influence of the acoustic streaming effects [6].

Computational fluid dynamic modelling was done in COMSOL Multiphysics, using microfluidic module as described in our previous study [7].

**Supplementary Figures**

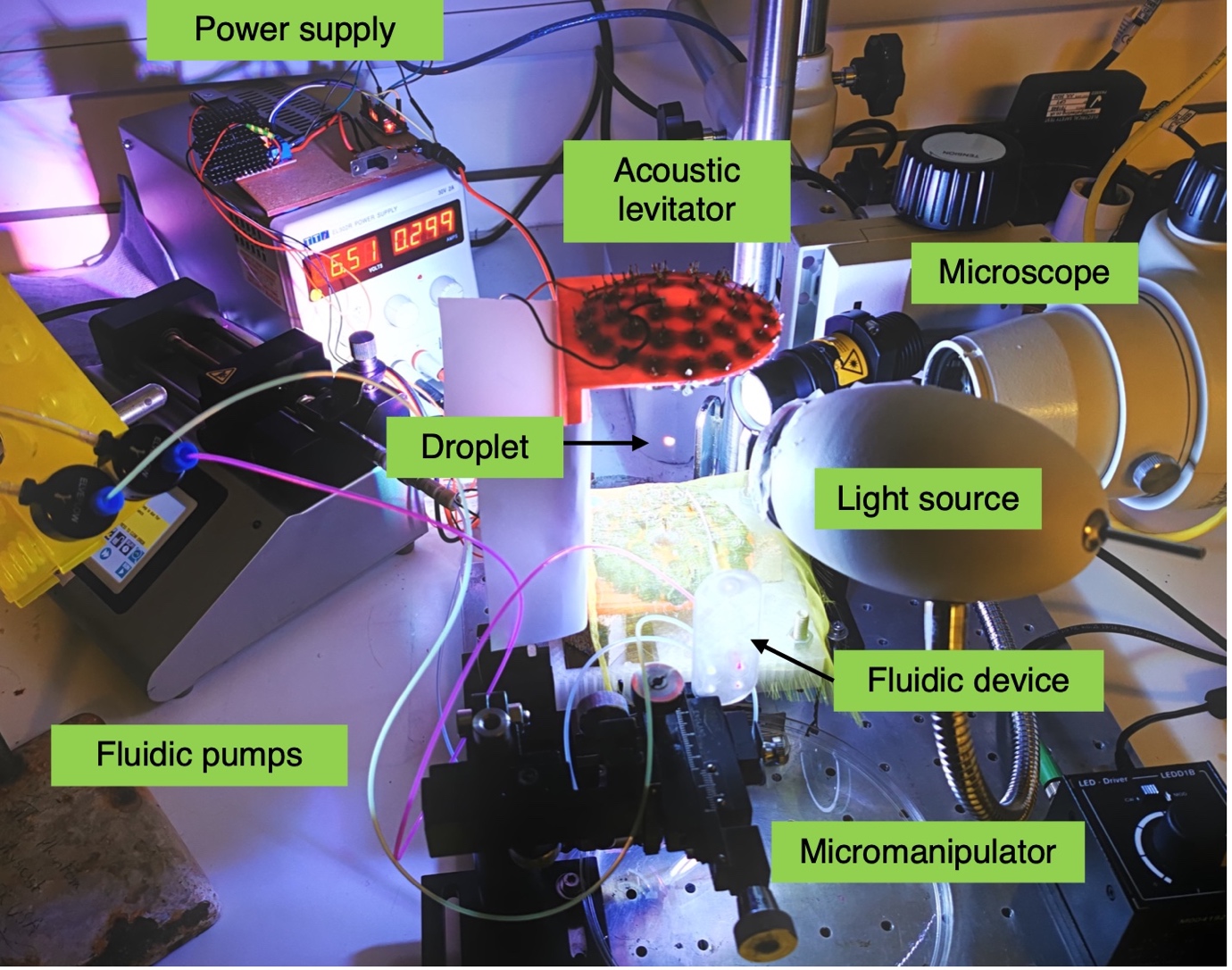

**Fig S1. Droplet laboratory experiment setup**. An ACDC droplet can be seen levitating in the acoustic trap.

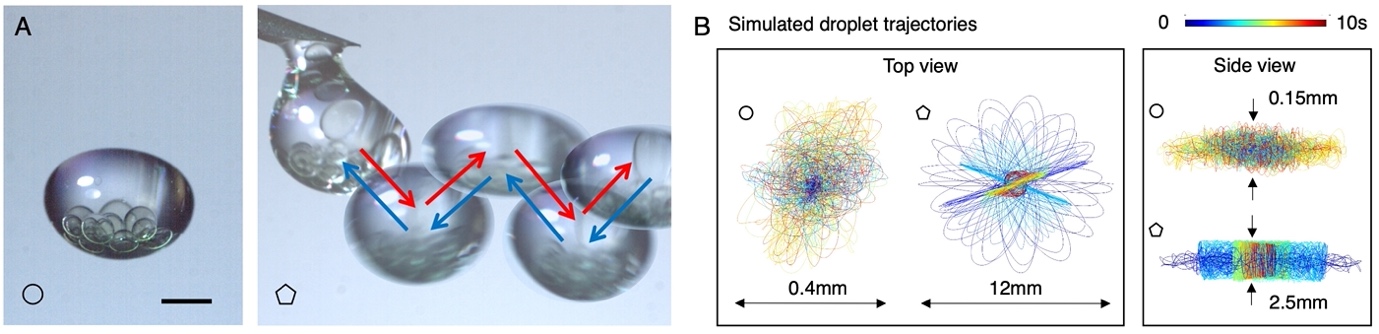

**Fig S2. Experimental and simulated droplet trajectories in the acoustic trap,** data complementary to Figure 1C. **A.** Left, an ACDC droplet stably levitated in air in the centre of an acoustic trapping node. Right, the trajectory (red followed by blue arrows) of a levitated ACDC droplet released from a position offset from the centre of the acoustic standing wave. Scale bar = 1 mm. **B.** Simulated trajectories of levitated droplets in comparison to that shown in Fig S2A. The trajectories marked with circles indicate that the droplet displays small oscillations both horizontally (< 200 µm) and vertically (<75 µm), even if levitated relatively stably in the acoustic standing wave. The trajectories marked with pentagons indicate the simulated trajectories when the droplet is released from an offset position as illustrated experimentally in Fig. S2A-right panel (pentagon labelled), where much larger oscillatory ranges are observed (6 mm).

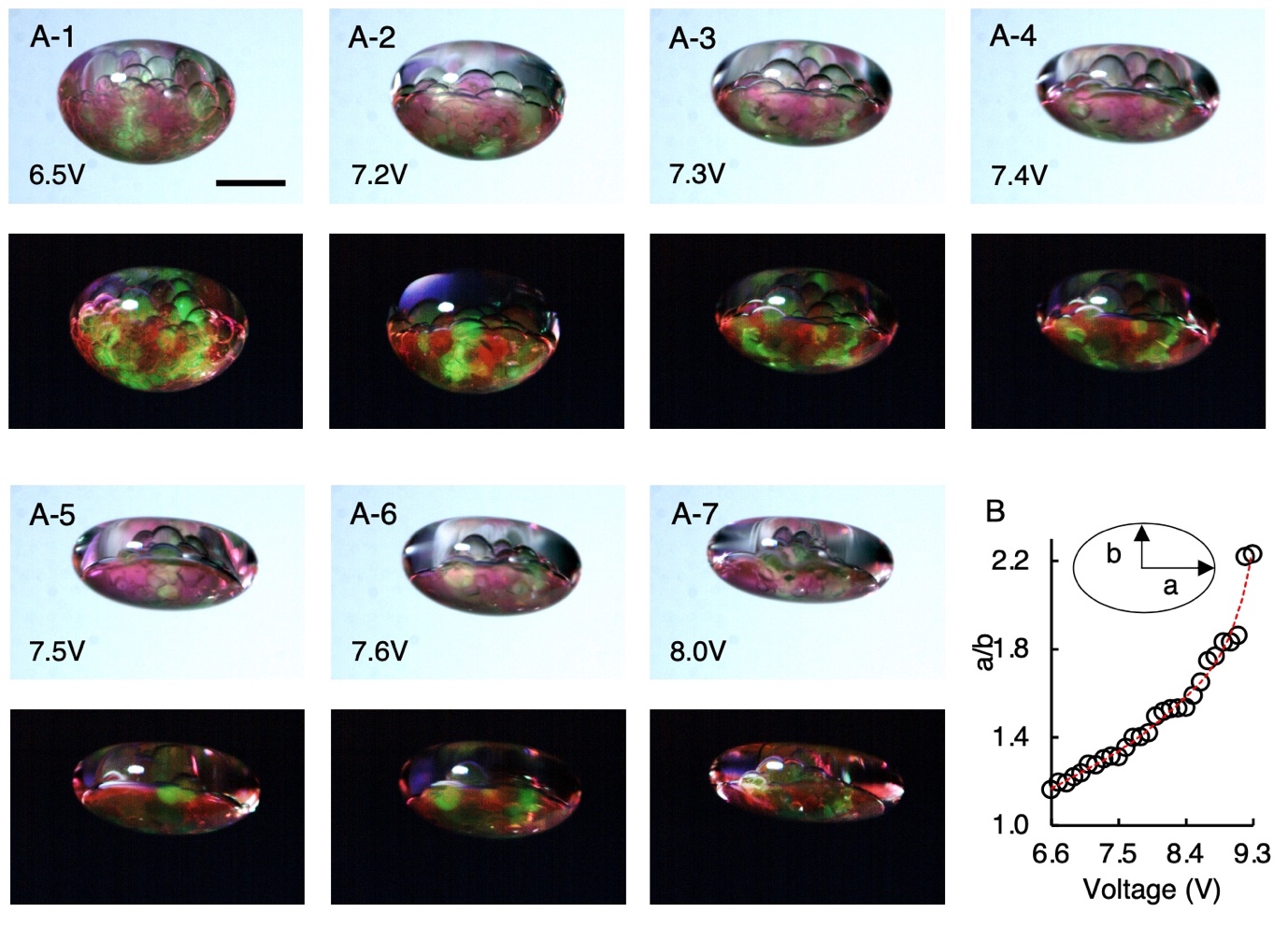

**Fig S3. The relationship between levitator input voltage and the levitated droplet shape. A1-7**. Example bright and dark images shows that the shape of a levitated ACDC droplet, which contained two types of cores (red and green), was gradually changed from ellipsoid to disk with progressive increases of input voltage and thus acoustic power to the levitator. Note that lipid bilayers remain stable throughout the shape manipulation with no coalescence of the cores. **B.** Measurement of the droplet aspect ratio (a/b) with increasing acoustic power (voltage to levitator) over the range 6.6. to 9.3 V. Representing quantitative analysis of the images of the type shown in A1-7.

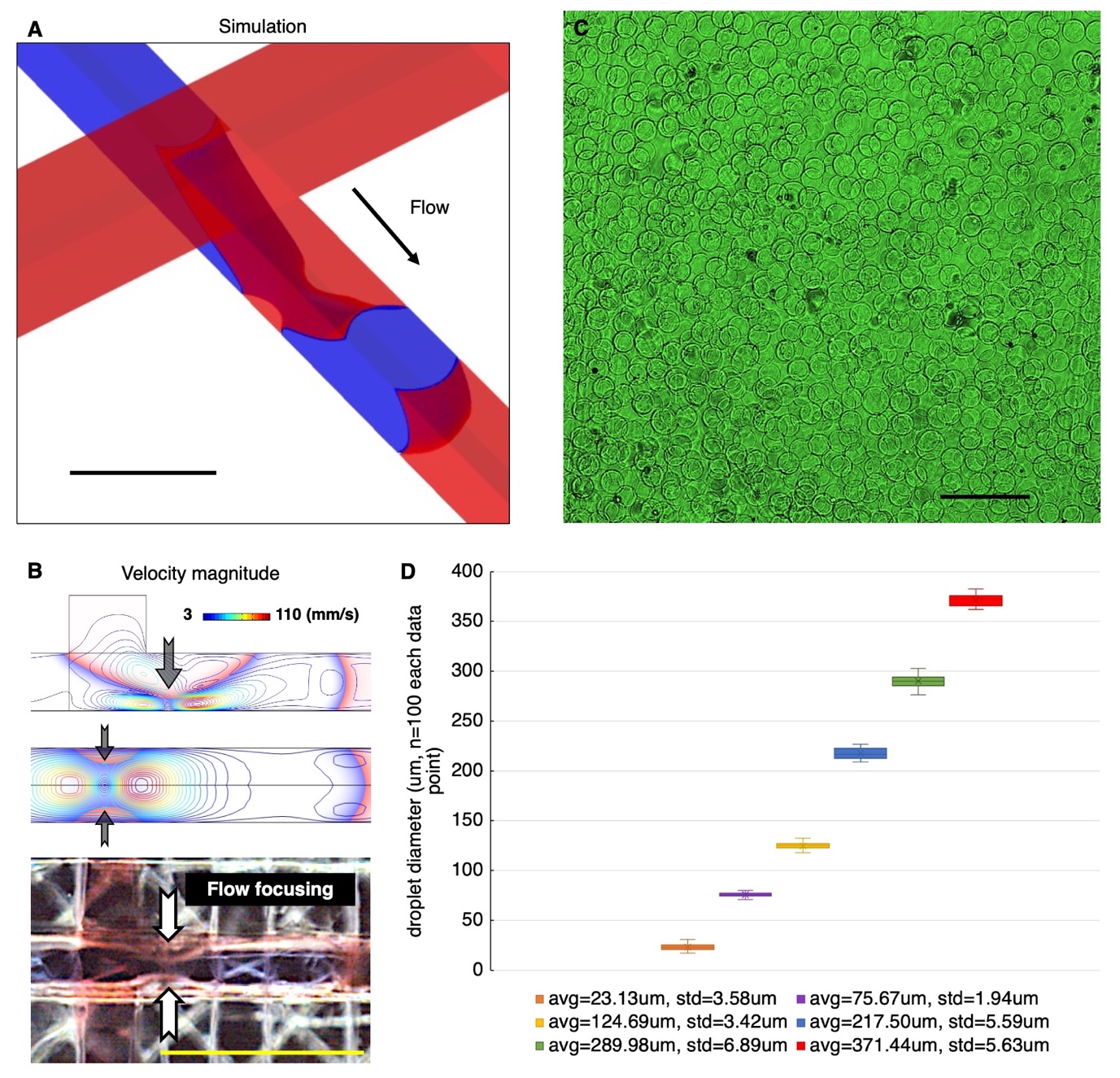

**Fig S4. Simulated and experimental details of 3D-printed, multi-layered, droplet forming fluidic junctions.** The multi-layered fluidic droplet forming junctions were designed to have the two input fluidic ducts stack on each other orthogonally, with a rectangular interface. This is to have sharp corners of the junction by shaping the interface with two straight printed lines, in comparison to the single layer junction that will naturally have roundish corners due to the movement resolution of the printhead. **A**. COMSOL simulation of water droplet (blue, 0.1 ml hr^-1^) at the point of break up in a continuous oil phase flow (red, 0.8 ml hr^-1^). Scale bar = 300 μm. **B.** Top, simulated velocity profiles during droplet breakup (side view and top view). The black arrows indicate the focusing flow. Bottom, corresponding image of water droplet (blue) break up in a mineral oil continuous phase flow (red) in a 3D-printed fluidic device. Scale bar = 0.5 mm. The visible cross-hatching effect is a consequence of the deposition lines of the polymer 3D printing process. **C.** A collection of alginate (average diameter = 124.69 μm) microgels formed by a multi-layered junction as illustrated in A and B. Scale bar = 1 mm. **D.** The uniformity of the droplet formed by the multi-layered junction, diameters ranging from 20 μm (approximate cell size) to 370 μm, which is controlled by the input flow rates ratio.

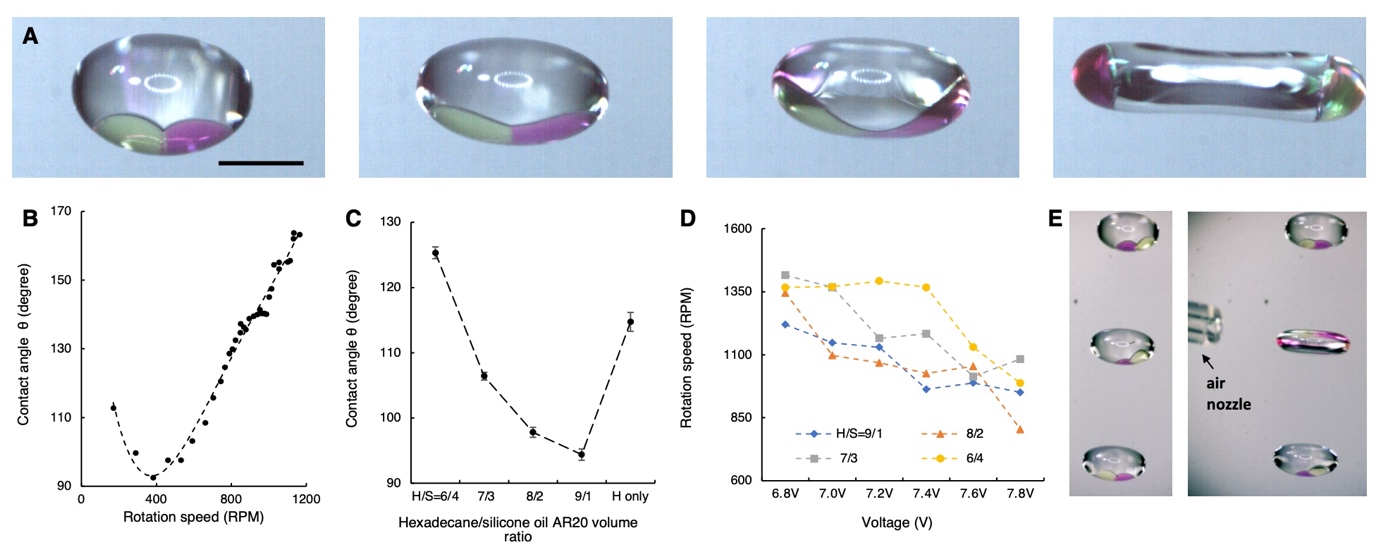

**Fig S5. Complementary data of pneumatic operations for the spinning of levitated ACDC droplets (Fig 3B). A,** From left to right, sequence of images showing aqueous cores (green and pink) connected by a lipid bilayer. Upon increasing spin speed due to increased air flow, the aqueous droplets move radially outwards in the less dense oil phase, reducing the bilayer area before eventually detaching. During the spinning action, the ACDC droplet shape progressively transforms from ellipsoid to a dumbbell shape, with the previously connected cores (green and pink) now spatially separated and residing at opposite ends of the dumbbell. Scale bar = 1 mm. **B,** The change of contact angle between the green and pink cores with increased spin rate of ACDC droplets. **C**, Contact angle of the red and green cores in different oil hexadecane (H) and silicone oil (S) mixtures. **D**, The spin rate of ACDC droplets under different applied voltages that gives rise to internal green and pink droplet detachment. Shown for different hexadecane and silicone oil mixtures. **E**. On levitation of multiple ACDC droplets in sequential nodes of the levitator, the pneumatic operation can be applied to a specific ACDC droplet without significantly influencing those levitating in neighbouring nodes of the levitator’s single axis acoustic standing wave field. Here a shape manipulation and droplet bilayer detachment is demonstrated on the middle ACDC droplet in a series of three.

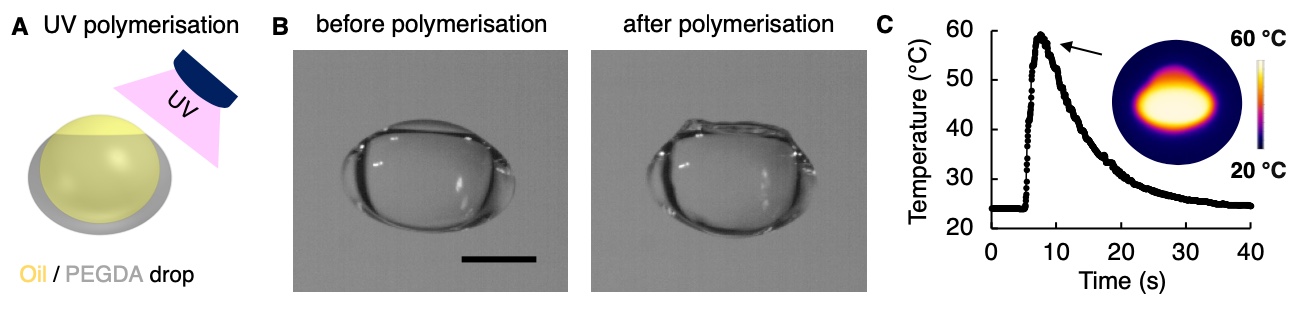

**Fig S6. Shell photo-polymerisation of a microfluidically-formed, levitated oil/PEGDA multi-phase droplet. A.** Schematic of the operation. **B.** images before and after the polymerisation. A bucket shaped solid shell was polymerised from the liquid PEGDA containing 1 wt% Irgacure 369 photoinitiator. Scale bar = 1 mm. **C**. The heat generated by the photopolymerisation and its dissipation during the reaction measured *in situ* using a thermal-imaging camera.

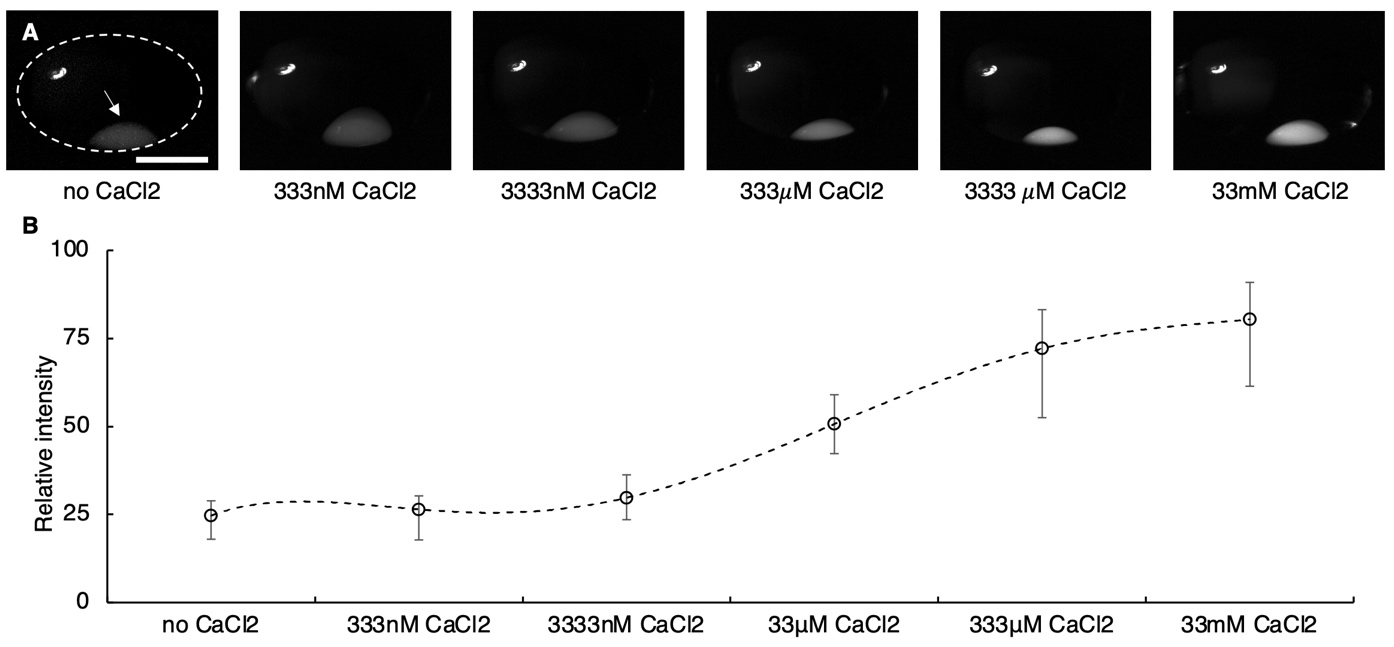

**Fig S7. Relative fluorescent intensity of the Fluo-8 containing cores (white) with different concentrations of pre-loaded CaCl_2_ in levitated ACDC droplets.** Scale bar denotes 1mm. The white dotted ellipsoid in panel 1 indicates the perimeter of the whole ACDC droplet. Droplet was not locked by magnetic manipulation.

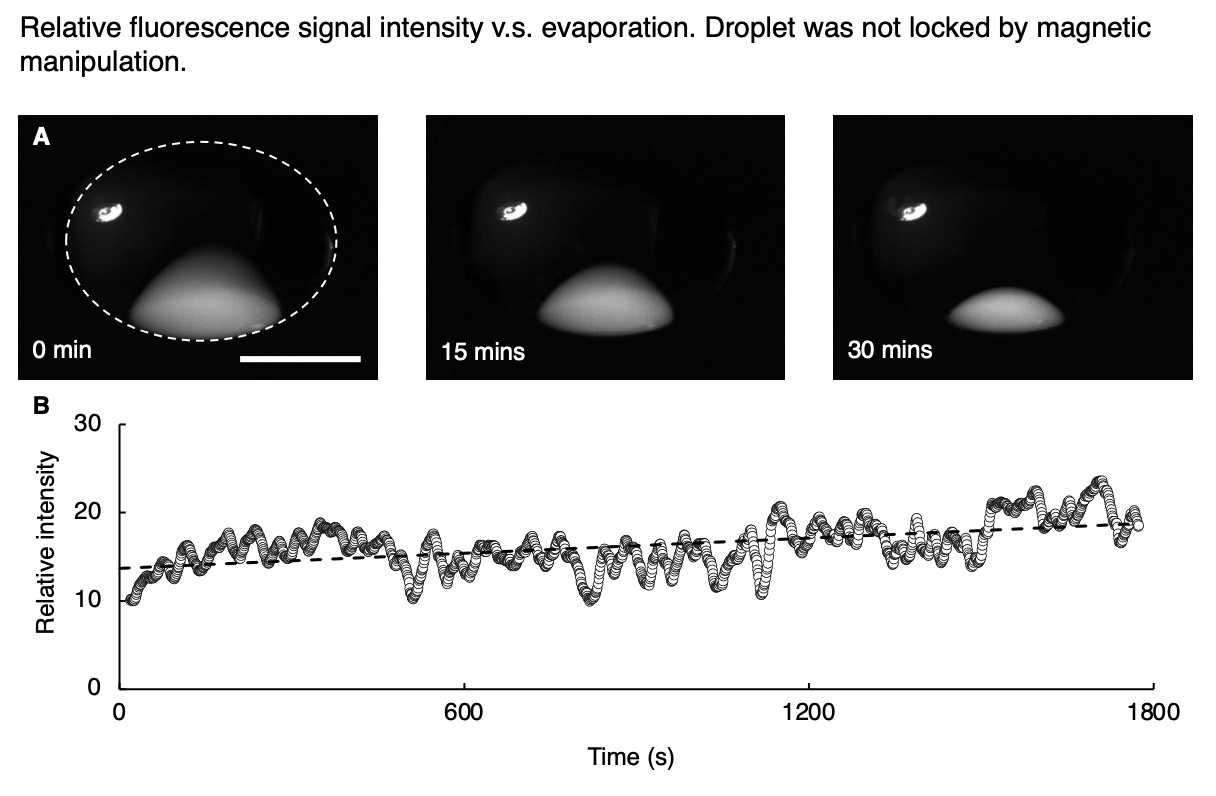

**Fig S8. Evaporation effects on fluorescence intensity are minimal compared to Ca^2+^ transport over the course of 30 minutes. The volume loss and the relative intensity change due to concentrating of the fluo-8 within an aqueous core (fluorescent white). Core also contains 3.3 mM CaCl_2_.** Scale bar = 1 mm. The white dotted ellipsoid indicates the edge of the whole ACDC droplet. Droplet was not locked by magnetic manipulation.

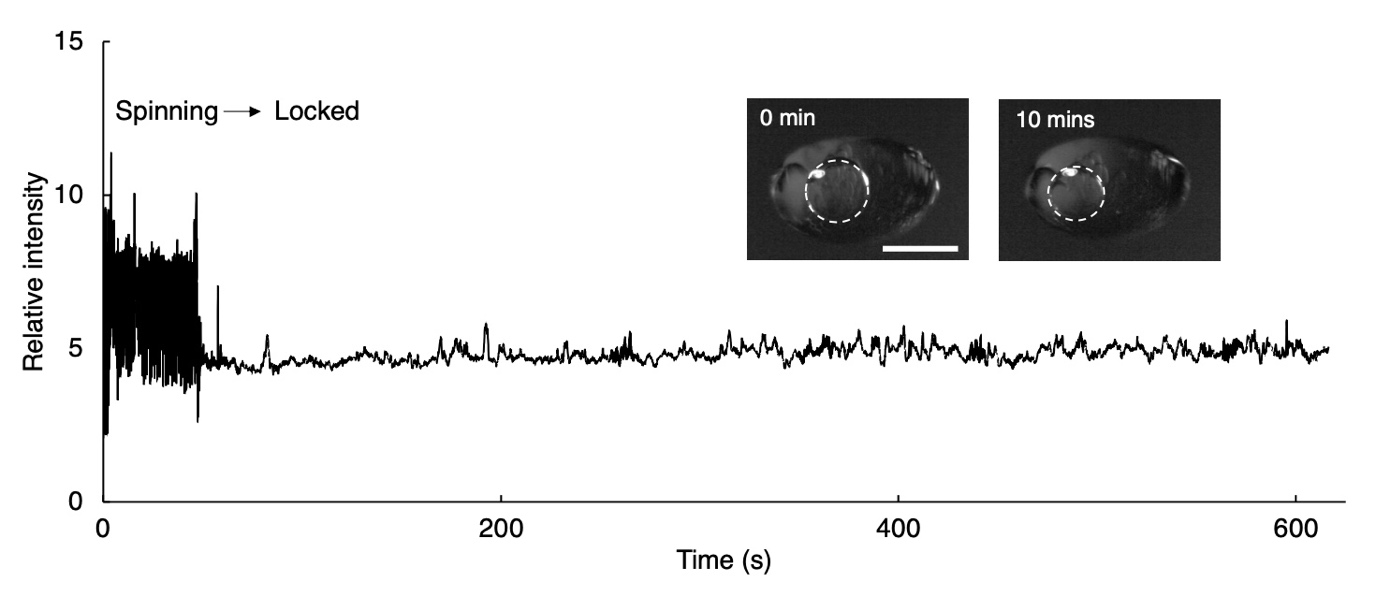

**Fig S9. Relative intensity of a control core (white dotted circle) containing fluo-8 but no MscL, in a levitated ACDC droplet.** The droplet was locked by magnetic manipulation. The lipid bilayers remain impermeable, with no significant intensity increase was observed. This result is complementary to Fig 5. The experimental droplet contains equivalent buffer materials to the MscL protein preparation, including a final concentration of 4 µM DDM, confirming that measured activation in MscL containing droplets (Fig4 and 5) is protein mediated.

**R**eference

1. Rosholm, K. R. et al. Activation of the mechanosensitive ion channel MscL by mechanical stimulation of supported Droplet-Hydrogel bilayers. *Scientific reports* **7,** 1-10 (2017).
2. Petrov, E., Rohde, P. R., & Martinac, B. Flying-patch patch-clamp study of G22E-MscL mutant under high hydrostatic pressure. *Biophysical journal* **100**, 1635-1641 (2011).
3. Marzo, A., Barnes, A., & Drinkwater, B. W.. TinyLev: A multi-emitter single-axis acoustic levitator. *Review of Scientific Instruments* **88**, 085105 (2017).
4. Schindelin J, et al. Fiji: an open-source platform for biological-image analysis. *Nat Methods* **9**, 676-82 (2012).
5. Bach, J. S., & Bruus, H. Theory of pressure acoustics with viscous boundary layers and streaming in curved elastic cavities. *The Journal of the Acoustical Society of America* **144**, 766-784 (2018).
6. Muller, P. B., Barnkob, R., Jensen, M. J. H., & Bruus, H. A numerical study of microparticle acoustophoresis driven by acoustic radiation forces and streaming-induced drag forces. *Lab on a Chip* **12**, 4617-4627 (2012).
7. Li, J., Lindley-Start, J., Porch, A., & Barrow, D. Continuous and scalable polymer capsule processing for inertial fusion energy target shell fabrication using droplet microfluidics. *Scientific reports* **7**, 1-10 (2017).
